# Developing a Multimodal miR-15a Mimic to Overcome Gemcitabine and RAS(ON) Inhibitor Daraxonrasib Resistance in Pancreatic Ductal Adenocarcinoma

**DOI:** 10.64898/2026.11.07.765071

**Authors:** Amartya Pal, Anushka Ojha, John Yuen, Hersh Bendale, Lei Chen, Iwao Ojima, Jingfang Ju

## Abstract

Gemcitabine remains a mainstay of pancreatic ductal adenocarcinoma (PDAC) treatment, while the RAS(ON) inhibitor daraxonrasib has recently demonstrated improved survival in previously treated metastatic disease. However, acquired resistance to both therapies remains a major clinical challenge. Here, we developed MTX-5-FU-Gem-miR-15a, a multimodal microRNA (miRNA)-based therapeutic, by chemically modifying tumor-suppressive miR-15a with methotrexate (MTX), 5-fluorouracil (5-FU), and gemcitabine (Gem) to combine sequence specific gene regulation with disruption of nucleotide metabolism. MTX-5-FU-Gem-miR-15a exhibited low-nanomolar potency across five PDAC cell lines and retained activity in gemcitabine- and daraxonrasib-resistant models under vehicle-free conditions, substantially exceeding that of stoichiometrically matched free antimetabolites. Treatment induced S-phase accumulation, increased DNA damage and apoptosis, and suppressed checkpoint and survival-associated proteins, including WEE1, CHK1, YAP1, and BCL-2. Transcriptomic and proteomic analyses revealed increased DNA replication, replication-stress, and repair-associated programs, together with reduced oncogenic signaling. In gemcitabine-resistant cells, treatment restored the gemcitabine-activating enzyme dCK, reduced ABCC1 and ABCC3 expression, and suppressed resistance-associated signaling programs, indicating partial remodeling of resistance-associated pathways. MTX-5-FU-Gem-miR-15a also synergized with daraxonrasib (mean Bliss synergy score, 22.52) and activated tumor-cell-intrinsic STING-TBK1 signaling, accompanied by increased interferon stimulated gene expression and IFN-β and CXCL10 secretion. Finally, MTX-5-FU-Gem-miR-15a retained low-nanomolar potency in three patient-derived PDAC organoids and significantly suppressed metastatic tumor growth *in vivo* at 4 mg/kg. Together, these findings support a multimodal miRNA therapeutic strategy that combines replication-associated DNA damage, checkpoint protein suppression, and innate immune activation-associated responses to target therapeutic resistance in PDAC.

## 1. Introduction

Pancreatic ductal adenocarcinoma (PDAC) remains one of the most lethal malignancies and is the third leading cause of cancer-related death in the United States. The five-year relative survival rate for pancreatic cancer is approximately 13%, declining to less than 5% in patients with metastatic disease [1]. Gemcitabine-based chemotherapy has been a mainstay of PDAC treatment for nearly three decades and is still widely used, particularly in combination with albumin-bound paclitaxel, while FOLFIRINOX (leucovorin, 5-fluorouracil, irinotecan, and oxaliplatin) provides another standard multi-agent chemotherapeutic regimen. Although these regimens improved outcomes, median overall survival in separate metastatic trials remained around 8.5 months with gemcitabine plus nab-paclitaxel and 11.1 months with FOLFIRINOX [2, 3]. More recently, daraxonrasib, a RAS(ON) multi-selective tri-complex inhibitor targeting the active GTP-bound forms of mutant and wild-type *KRAS*, *NRAS*, and *HRAS*, has shown meaningful activity in previously treated RAS-mutant metastatic PDAC and doubled the overall survival of PDAC patients relative to chemotherapy in a randomized phase 3 trial [4, 5]. Despite their distinct mechanisms of action, acquired resistance remains an important limitation of both cytotoxic chemotherapy and RAS-directed targeted therapy [6, 7]. Therefore, understanding and overcoming these resistance mechanisms is critical to extending the benefit of both established and emerging therapies.

Gemcitabine resistance in PDAC arises through several mechanisms involving drug transport, metabolism, and adaptive survival signaling. Reduced expression of the nucleoside transporter hENT1 (SLC29A1) or the activating enzyme deoxycytidine kinase (dCK), together with increased cytidine deaminase (CDA) activity and elevated ribonucleotide reductase subunits RRM1 and RRM2, can substantially reduce gemcitabine activity [8, 9]. Consistent with this, low hENT1 and dCK expression have also been associated with poorer outcomes in patients receiving gemcitabine-based therapy [10–12]. Beyond altered drug metabolism, resistant PDAC cells frequently acquire mesenchymal and stem-like features and become increasingly dependent on DNA damage checkpoint signaling to survive persistent replication stress [13–16]. Together, these changes and PDAC’s dense, poorly perfused stromal microenvironment create multiple overlapping mechanisms that sustain gemcitabine resistance [17].

Resistance to RAS(ON) inhibition develops through a distinct mechanism. In patients progressing on daraxonrasib, acquired alterations largely converged on reactivation of RAS pathway signaling, most commonly through amplification of mutant *KRAS* together with alterations in RTK, MAPK, and PI3K signaling [6]. Mutant *KRAS* amplification was strongly associated with pretreatment TP53 loss, an alteration present in roughly 70% of PDAC, and resistant cells carrying this alteration showed increased dependence on DNA damage response pathways, including the ATR/CHK1/WEE1 checkpoint axis [6, 18]. Although inhibiting this pathway can suppress resistant cells, the toxicity of direct checkpoint inhibitors has limited their clinical use [19, 20]. Together, these findings point to a broader challenge in PDAC: interconnected metabolic, oncogenic, stemness, and checkpoint-survival programs sustain resistance and compensate for one another when any single mechanism is inhibited, favoring approaches that disrupt several simultaneously.

MicroRNAs (miRNAs) are particularly attractive therapeutic candidates to address this complexity because a single miRNA can regulate multiple genes across interconnected signaling pathways [21]. Through partial sequence complementarity with target mRNAs, miRNAs can suppress several resistance-associated programs simultaneously, offering an advantage over therapies that inhibit only one pathway at a time [22, 23]. However, despite this strong biological rationale, no miRNA-based cancer therapeutic has been FDA-approved, largely because instability, inefficient delivery, and limited tumor specificity have hindered clinical translation [24].

Previous studies from our group have identified hsa-miR-15a as a tumor suppressor with promising therapeutic relevance in PDAC. Mature miR-15a is reduced in PDAC tissues and low expression is associated with reduced overall survival (high versus low miR-15a, HR = 0.47, 95% CI 0.30-0.72; log-rank p = 0.00043; Fig. 1A), while the co-transcribed miR-16 remains largely unchanged, suggesting impaired miR-15a processing in PDAC rather than loss of the miR-15a/16-1 locus at chromosome 13q14 [22]. We also identified WEE1 and CHK1 as direct targets of miR-15a in PDAC, with two miR-15a binding sites in each transcript validated by reporter assays and by suppression at both the mRNA and protein levels [22, 25]. Other oncogenic targets, including BMI1 and YAP1, are also suppressed by miR-15a. These targets are closely linked to the resistance mechanisms examined in this study. WEE1 and CHK1 maintain the DNA damage checkpoint and help cells survive gemcitabine-induced replication stress, and dependence on this checkpoint axis has also been identified in RAS(ON) inhibitor-resistant cells [6, 14–16, 22]. BMI1 promotes self-renewal and EMT and has been associated with gemcitabine resistance, whereas YAP1 can support tumor-cell survival independently of oncogenic *KRAS* and contributes to escape from *KRAS* dependence [26, 27]. Targeting these pathways individually has shown limited activity in PDAC. The CHK1 inhibitor LY2603618 failed to improve the efficacy of gemcitabine in a phase II trial, and WEE1 inhibition alone did not produce tumor regression [19, 28, 29]. Restoring tumor suppressor miR-15a therefore offers a unique approach, allowing a single regulatory program to suppress multiple checkpoint, survival, and resistance-associated pathways.

**Figure 1.**
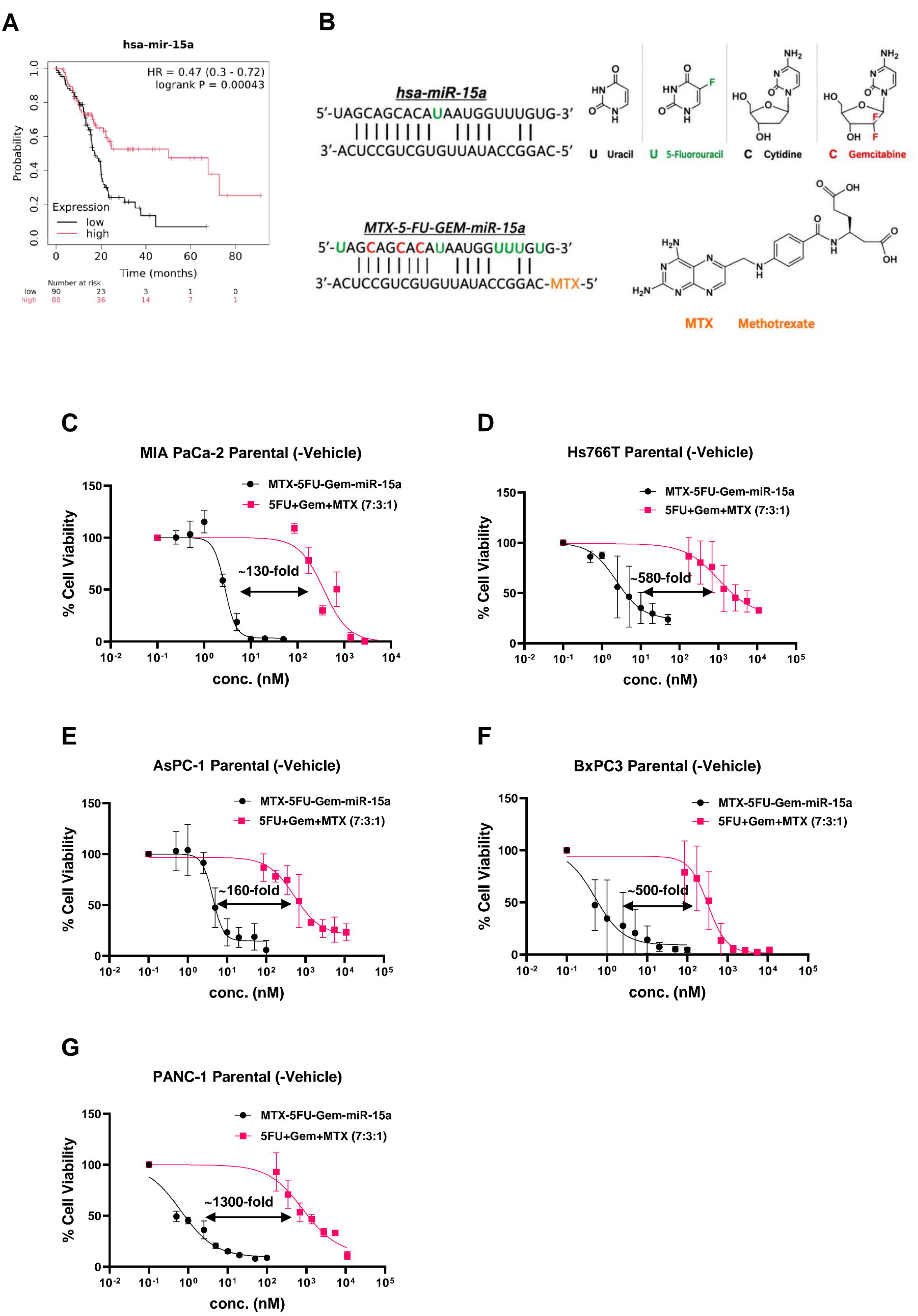
Design of MTX-5-FU-Gem-miR-15a and cytotoxic activity across pancreatic cancer cell lines. (A) Kaplan–Meier survival analysis showing the association between miR-15a expression and overall survival in patients with PDAC. (B) Schematic representation of hsa-miR-15a and MTX-5-FU-Gem-miR-15a, showing substitution of uridine and cytidine residues in the miR-15a guide strand with 5-fluorouracil (5-FU) and gemcitabine (Gem), respectively, and conjugation of methotrexate (MTX) to the passenger strand. (C–G) Dose-response curves showing cell viability following vehicle-free treatment with MTX-5-FU-Gem-miR-15a or a stoichiometrically matched free-drug combination of 5-FU, Gem, and MTX (7:3:1) in parental MIA PaCa-2 (C), Hs 766T (D), AsPC-1 (E), BxPC-3 (F), and PANC-1 (G) cells. Cell viability was assessed after six days of treatment. Data represent mean ± SD from n = 3 biological replicates.

To realize the therapeutic potential of miR-15a, we have designed a multimodal miR-15a mimic to overcome the major barriers of nucleic acid-based therapy by enhancing stability, therapeutic efficacy, and ease of delivery. Specifically, MTX-5-FU-Gem-miR-15a was synthesized in which uridine (U) and cytidine (C) residues within the miR-15a guide strand were substituted with 5-fluorouracil (5-FU) and gemcitabine (Gem), respectively, while folate analog methotrexate (MTX) was conjugated to the passenger strand (Fig. 1B). These modifications combine the tumor suppressor function of miR-15a with three antimetabolite activities that converge on nucleotide metabolism and DNA replication through distinct mechanisms, as MTX-5-FU-Gem-miR-15a will release 5-FU, Gem and MTX once degraded by both 5’- and 3’-exonucleases. 5-FU inhibits thymidylate synthase, depleting dTMP and causing DNA damage; gemcitabine causes DNA chain termination and inhibits ribonucleotide reductase; and MTX inhibits dihydrofolate reductase, restricting the reduced folate pool required for thymidylate and de novo purine synthesis [30–33]. Simultaneous interference with nucleotide metabolism at multiple points is difficult for tumor cells to compensate for and is particularly relevant to gemcitabine resistance, where expansion of competing deoxynucleotide pools represents an important mechanism by which resistant cells reduce gemcitabine activity [7, 9].

MTX provides additional functions beyond its effects on folate metabolism. At low doses, MTX has been reported to activate cGAS-STING signaling and increase type I interferon responses, providing a potential immunomodulatory component to the construct [34]. This is particularly relevant in PDAC, where a poorly immunogenic tumor microenvironment contributes to limited responses to immune checkpoint therapy [35]. MTX also binds folate receptor alpha (FRα) with high affinity, providing an additional potential mechanism for tumor-directed uptake in FRα-expressing cells [36, 37].

MTX-5-FU-Gem-miR-15a is designed to function as a single molecular platform rather than as a mixture of separately administered drugs. By incorporating 5-FU and gemcitabine directly into the miR-15a guide strand and conjugating MTX to the passenger strand, the construct couples miR-15a-mediated target suppression with simultaneous disruption of nucleotide metabolism and DNA replication. This coordinated design also avoids the pharmacologic separation and asynchronous exposure that can occur when drugs are combined independently. Importantly, the modified miRNA remains active without a delivery vehicle, addressing one of the major barriers to therapeutic miRNA delivery. Incorporation of gemcitabine into a construct intended for gemcitabine-resistant disease is also mechanistically relevant. Because the gemcitabine moiety is delivered as part of the oligonucleotide scaffold, the cellular entry of MTX-5-FU-Gem-miR-15a does not require the nucleoside transport pathway needed for free gemcitabine, providing a way to test whether this platform can retain activity after resistance to conventional gemcitabine develops. In addition, both 5-FU and gemcitabine are established components of PDAC therapy, further supporting their incorporation into a therapeutic designed for this disease.

In this study, we investigated the therapeutic efficacy of MTX-5-FU-Gem-miR-15a in both treatment-naïve and drug-resistant PDAC models. We generated gemcitabine-resistant PDAC derivatives of MIA PaCa-2 and Hs 766T, in which the gemcitabine IC_50_ increased by more than 25-fold, and separately, a daraxonrasib-resistant MIA PaCa-2 model in which the daraxonrasib IC_50_ increased by more than 50-fold. MTX-5-FU-Gem-miR-15a showed low nanomolar potency across five PDAC cell lines and in all three resistant derivatives under vehicle-free conditions and retained activity in three-dimensional patient-derived organoid models. Transcriptomic profiling defined resistance-associated programs in gemcitabine-resistant cells and determined how treatment altered these pathways. Treatment opposed a substantial fraction of the resistance-associated transcriptional program and induced an interferon-stimulated gene program in both parental and resistant cells. Functionally, MTX-5-FU-Gem-miR-15a showed synergistic activity in combination with daraxonrasib within a defined concentration range, and suppressed tumor growth *in vivo*. Together, these findings support MTX-5-FU-Gem-miR-15a as a multimodal miRNA therapeutic that targets oncogenic and metabolic signals and disrupts checkpoint and survival pathways that sustain drug-resistant PDAC across mechanistically distinct resistance states.

## 2. Materials and methods

### 2.1 Cell lines and culture conditions

The human PDAC cell lines MIA PaCa-2 (RRID:CVCL_0428), PANC-1 (RRID:CVCL_0480), BxPC-3 (RRID:CVCL_0186), AsPC-1 (RRID:CVCL_0152), and Hs 766T (RRID:CVCL_0334) were obtained from ATCC. These cell lines were selected to represent the genetic and phenotypic heterogeneity of PDAC, including *KRAS* G12C (MIA PaCa-2), G12D (PANC-1 and AsPC-1), Q61H (Hs 766T), and *KRAS*-wild-type (BxPC-3) backgrounds. The panel also included cell lines with distinct epithelial and mesenchymal-like phenotypes associated with classical and basal-like PDAC molecular subtypes. MIA PaCa-2 and PANC-1 have been characterized as basal-like/mesenchymal models, while Hs 766T exhibits a mesenchymal phenotype. In contrast, AsPC-1 and BxPC-3 have been classified as non-mesenchymal, with features associated with the classical subtype [38]. Cells were maintained at 37°C in a humidified incubator with 5% CO₂. MIA PaCa-2, PANC-1, and Hs 766T cells were cultured in DMEM supplemented with 10% fetal bovine serum (FBS), whereas BxPC-3 and AsPC-1 cells were cultured in RPMI-1640 supplemented with 10% FBS. FBS was filter-sterilized using a Steriflip-GP 0.22 µm filter unit. All cell lines were authenticated by short tandem repeat profiling and tested negative for Mycoplasma contamination.

### 2.2 Generation of gemcitabine- and daraxonrasib-resistant cell lines

Gemcitabine-resistant derivatives were generated from MIA PaCa-2 and Hs 766T cells, which differ in growth kinetics and baseline gemcitabine sensitivity. Cells were exposed to stepwise increasing concentrations of gemcitabine (0.5 to 50 µM for MIA PaCa-2; 1 to 50 µM for Hs 766T), with each escalation applied only after cells resumed normal proliferation at the preceding dose. Selection was carried out over 3 to 4 months until stable resistant populations were established, designated MIA PaCa-2/GemR and Hs 766T/GemR.

A daraxonrasib-resistant derivative was generated in parallel from MIA PaCa-2 cells by continuous exposure to increasing concentrations of daraxonrasib (RMC-6236) from 10 to 500 nM over 3 months, designated MIA PaCa-2/DaraxR. This model was included because RAS(ON) multi-selective inhibition is now an emerging standard in previously treated metastatic PDAC, and acquired resistance to this class represents a distinct and clinically imminent problem.

Following drug selection, resistant derivatives were cultured in drug-free medium for two weeks before experimental treatment. Resistance was reassessed by comparing half-maximal inhibitory concentration (IC₅₀) values between parental and resistant derivatives following drug withdrawal to determine whether the reduced drug sensitivity persisted in the absence of continued selection pressure.

Hs 766T-Luc cells were generated by transducing parental Hs 766T cells with a luciferase-expressing lentiviral construct. Transduced cells were selected using puromycin, and luciferase expression was confirmed by *In vivo* Imaging System (IVIS) prior to use *in vivo*.

### 2.3 Three-dimensional culture

hF3, hF44, and hT89 organoid cells were provided by the Stony Brook Medicine Biobank (Stony Brook Medicine Biobank, Renaissance School of Medicine, Stony Brook, USA). Organoid cells were cultured and treated as previously described [25]. In brief, organoid cells were cultured with Corning Matrigel GFR Membrane Matrix (Corning Incorporated, Corning, NY, USA) and incubated at 37°C in Human Complete Feeding Medium (hCPLT) consisting of the following: Advanced DMEM/F12 (Thermo Fisher Scientific, Waltham, MA, USA), HEPES (final concentration, 10 mmol/L) (Thermo Fisher Scientific, Waltham, MA, USA), GlutaMAX (final concentration, 1x) (Thermo Fisher Scientific, Waltham, MA, USA), A 83-01 (final concentration, 500 nmol/L) (Tocris Bioscience, Bristol, UK), hEGF (final concentration, 50 ng/mL) (Thermo Fisher Scientific, Waltham, MA, USA), mNoggin (final concentration, 100 ng/mL) (Thermo Fisher Scientific, Waltham, MA, USA), hFGF10 (final concentration, 100 ng/mL) (Thermo Fisher Scientific, Waltham, MA, USA), hGastrin I (final concentration, 0.01 μmol/L) (Tocris Bioscience, Bristol, UK), N-acetylcysteine (final concentration, 1.25 mmol/L) (MilliporeSigma, St. Louis, MO, USA), nicotinamide (final concentration, 10 mmol/L) (MilliporeSigma, St. Louis, MO, USA), prostaglandin E2 (final concentration, 1 μmol/L) (Tocris Bioscience, Bristol, UK) B27 supplement (final concentration, 1X) (Thermo Fisher Scientific, Waltham, MA, USA), R-spondin1-Conditioned Media (final concentration, 10%) (Thermo Fisher Scientific, Waltham, MA, USA), and afamin/Wnt3A conditioned media.

### 2.4 Design and synthesis of MTX-5-FU-Gem-miR-15a and control constructs

In MTX-5-FU-Gem-miR-15a, the guide strand was modified by substituting uridine and cytidine residues with 5-FU and Gem. The passenger strand was functionalized with a 5′ azide linker and conjugated with methotrexate (MTX) via dibenzocyclooctyne (DBCO)-mediated copper-free click chemistry. Modified oligonucleotides were synthesized and HPLC-purified (Horizon Discovery) and MTX-DBCO was synthesized at Stony Brook University. The MTX-linked passenger strand was purified using the Oligo Clean & Concentrator Kit (Zymo Research) and annealed with the guide strand at 60°C to generate the final duplex. The miR-15a guide strand contains seven uridine and three cytidine residues; complete substitution therefore incorporates seven 5-FU and three Gem nucleoside analogs per guide strand. A single MTX molecule is conjugated to the passenger strand, resulting in a fixed 7:3:1 stoichiometry of 5-FU:Gem:MTX within each MTX-5-FU-Gem-miR-15a duplex.

### 2.5 MicroRNA transfection and drug treatment

For vehicle-mediated transfection, 1.5 × 10 cells per well were seeded in 6-well plates and transfected 24 hours later with 50 nM of MTX-5-FU-Gem-miR-15a, unmodified miR-15a, or negative control using Oligofectamine (Thermo Fisher Scientific). Five hours post-transfection, the medium was replaced with fresh medium supplemented with 10% dialyzed FBS (DFBS). For vehicle-free delivery, cells were seeded at 1,500 cells per well in 96-well plates and treated directly with the respective oligonucleotides or drug controls diluted in culture medium. After 24 hours, the medium was replaced with fresh medium containing 10% DFBS.

Free-drug controls were prepared at molar equivalents matched to the stoichiometry of the construct. The miR-15a guide strand contains seven uridine and three cytidine residues, each substituted with 5-FU and Gem respectively, and one MTX is conjugated per passenger strand; each duplex therefore carries 5-FU, Gem, and MTX in a fixed 7:3:1 ratio. Accordingly, individual free-drug controls were tested at matched component ratios of 5-FU and Gem (7:3) or 5-FU, Gem and MTX (7:3:1) to evaluate whether individual drug combinations could recapitulate the activity of the miRNA mimic.

### 2.6 Cell viability and cytotoxicity assays

Cells were seeded in 96-well plates at 1500 cells/well concentration and treated with increasing concentrations of miRNA constructs, gemcitabine, daraxonrasib, or free-drug controls after 24 hours. After six days of treatment, viability was measured using the WST-1 reagent (MilliporeSigma), with absorbance read at 450 nm against a 630 nm reference.

Organoid cells were dissociated into single cell suspension using TrypLE and plated at 500 cells/ well in 50 μL of 10% Matrigel/ hCPLT in black, clear bottom 96-well plate. 72-hours post-plating and visual confirmation of organoid formation, organoids were treated with either MTX-5-FU-Gem-miR-15a or Gemcitabine and after 6 days, cell viability was examined using CellTiter-Glo 3D (Promega).

Relative cell viability was calculated by normalizing absorbance or luminescence values to the corresponding negative-control cells.

### 2.7 Cell cycle and apoptosis analysis

For cell-cycle analysis, treated cells were harvested after 72 hours, resuspended in Krishan-modified buffer containing RNase H and propidium iodide, and analyzed on a CytoFLEX and LSRFortessa flow cytometer. Cell-cycle distribution was quantified using ModFit LT software and is reported as the fold change in G2/S ratio relative to negative control. For apoptosis assays, cells were stained with Annexin V-FITC and propidium iodide 72 hours after treatment, and Annexin V-positive cells (early plus late apoptotic) were quantified by flow cytometry. Fold change in apoptotic cells was calculated relative to negative-control cells.

### 2.8 Western blot analysis

PDAC cells were treated or transfected with the indicated constructs or controls as described above. After 72 h, cells were lysed in RIPA buffer (MilliporeSigma) supplemented with protease and phosphatase inhibitor cocktail (MilliporeSigma), and protein concentrations were determined by Bradford assay. Equal amounts of protein were separated by SDS-PAGE and transferred to PVDF membranes.

Membranes were blocked and probed with the following primary antibodies: rabbit anti-WEE1 (Cell Signaling Technology (CST), 13084, 1:1,000), mouse anti-CHK1 (CST, 2360, 1:500), rabbit anti-YAP1 (CST, 4912, 1:1,000), rabbit anti-BMI1 (CST, 6964, 1:1,000), mouse anti-BCL-2 (CST, 15071, 1:1,000), rabbit anti-BAX (CST, 2772, 1:1,000), rabbit anti-BAK (CST, 12105, 1:1,000), rabbit anti-CCND1 (CST, 55506, 1:1,000), mouse anti-p21 Waf1/Cip1 (CST, 2946, 1:2,000), rabbit anti-GADD45α (CST, 4632, 1:1,000), rabbit anti-phospho-Histone H2A.X/γH2AX Ser139 (CST, 2577, 1:1,000), rabbit anti-RAD51 (CST, 8875, 1:1,000), rabbit anti-PARP (CST, 9532, 1:1,000), rabbit anti-cleaved PARP Asp214 (CST, 5625, 1:1,000), rabbit anti-caspase-7 (CST, 12827, 1:1,000), rabbit anti-cleaved caspase-7 (CST, 8438, 1:1,000), rabbit anti-phospho-STING Ser366 (CST, 19781, 1:1,000), rabbit anti-phospho-TBK1 Ser172 (CST, 5483, 1:1,000), anti-MRP1 (Proteintech, 67228-1, 1:1,000), anti-MRP3 (Proteintech, 25358-1, 1:1,000), anti-dCK (Proteintech, 17758-1, 1:1,000), anti-CDA (Abcam, ab222515, 1:500) and mouse anti-β-actin (MilliporeSigma, A5441, 1:10,000) and β-tubulin (CST, 2146, 1:5,000).

Primary antibodies were diluted in 5% milk or 5% bovine serum albumin in TBST according to antibody-specific recommendations, and membranes were subsequently probed with goat anti-rabbit IgG-HRP (Bio-Rad, 172-1019, 1:5,000) or goat anti-mouse IgG-HRP (Bio-Rad, 1706516, 1:5,000). Bands were visualized on a LI-COR Biosciences Odyssey Fc imaging system after incubation with SuperSignal West Pico PLUS Chemiluminescent Substrate (Thermo Fisher Scientific) and quantified using Image Studio software version 5.2.4 (LI-COR Biosciences). For each target, expression was normalized to β-actin or α-tubulin and reported relative to the negative-control sample, which was set to 1. Densitometry is reported as the mean of n = 3 independent biological replicates.

### 2.9 RNA-seq Analysis

MIA PaCa-2 parental (PR) and gemcitabine-resistant (GR) cells were treated with negative control (NC) or M15a (MTX-5-FU-Gem-miR-15a) at 50 nM dose as described above, and total RNA was isolated 72 h after treatment using TRIzol reagent (Thermo Fisher Scientific). Poly(A)-enriched mRNA sequencing and initial raw-data processing were performed by Novogene. The analysis included four groups (PR NC, PR M15a, GR NC, and GR M15a) with three biological replicates per group.

Gene-level differential expression analysis was performed in R using DESeq2 [39]. Genes were retained when they had at least 10 counts in at least three samples. The principal comparisons included acquired resistance (GR NC vs PR NC), M15a treatment in parental cells (PR M15a vs PR NC), and MTX-5-FU-Gem-miR-15a treatment in resistant cells (GR M15a vs GR NC). For ranked pathway analyses, genes were ordered using the unshrunken DESeq2 Wald statistic.

Gene set enrichment analysis was performed using clusterProfiler and fgsea with curated KEGG, Hallmark, and Reactome gene sets [40–43]. Pathway analyses focused on cancer-associated signaling, nucleotide metabolism, DNA replication and repair, cell-cycle/checkpoint regulation, apoptosis, drug-resistance programs, and innate immune signaling. Selected pathway modules were also summarized using gene-level expression scores to compare DNA-damage, replication-stress, checkpoint, mitotic, and interferon-associated programs across the four experimental conditions.

Principal component analysis was performed using variance-stabilized expression data to evaluate separation among experimental groups. For heatmap visualization, normalized gene expression values were transformed into row-wise Z-scores across individual samples and then averaged within each experimental condition. Curated heatmaps were generated for replication-stress genes, cell-cycle and mitotic regulators, checkpoint-associated genes, ABC transporters, apoptosis/survival genes, and interferon-stimulated genes. To quantify reversal of resistance-associated pathway activity, a pathway-reversion score was calculated as follows:
Reversion score = −(NES_resistance × NES_M15a in GemR)
where NES_resistance was obtained from GemR NC versus PAR NC and NES_M15a in GemR from GemR M15a versus GemR NC. Positive scores indicate pathways whose resistance-associated direction was opposed by MTX-5-FU-Gem-miR-15a treatment.

### 2.10 Proteomic analysis

MIA PaCa-2 parental (PR) cells were treated with negative control (NC) or MTX-5-FU-Gem-miR-15a at 50 nM dose as described above, and total cell lysate was extracted 72 h after treatment using 5% SDS supplemented with protease and phosphatase inhibitor cocktail (MilliporeSigma). The samples were processed and analyzed using Mass Spectrometry by the Proteomics Core at Stony Brook University.

Quantitative proteomic data from M15a-treated and control cells were analyzed in R using normalized protein-abundance measurements from two M15a and two NC samples. Only proteins with high identification confidence, at least one unique peptide, and complete quantitative measurements across all four samples were included in the primary ranked analysis. Missing abundance values were not imputed as zero.

Protein abundances were log₂ transformed, and differential protein abundance was evaluated using limma, with proteins ranked by the moderated *t*-statistic for M15a versus NC [44]. Gene set enrichment analysis was then performed to identify coordinated changes in pathways associated with nucleotide metabolism, DNA replication and repair, cell-cycle progression, mitotic regulation, apoptosis, oncogenic signaling, and innate immune responses.

For visualization of selected protein programs, protein abundances were converted to row-wise Z-scores across the individual samples and averaged within the NC and M15a groups. Curated heatmaps were generated for proteins involved in G2/M and mitotic execution, apoptosis, and innate/interferon-associated signaling. Leading-edge proteins contributing to selected enriched pathways were additionally examined to identify proteins driving the observed pathway-level changes.

### 2.11 Combination and synergy analysis

MTX-5-FU-Gem-miR-15a and daraxonrasib were tested across an 8 × 6 concentration matrix covering 0 to 10 nM of MTX-5FU-gemmir15a and 0 to 500 nM of daraxonrasib in MIA Paca2/daraxR cells. Cells were seeded in 96-well plates and 24 hours later, treated with drug combination, and viability was measured after 6 days as described above. Drug interactions were quantified using the Bliss independence model in SynergyFinder [45]. Mean Bliss synergy scores greater than 10 were considered synergistic, and scores between 0 and 10 were considered additive; the concentration region over which synergy was observed is reported alongside the global score, since a positive global score can arise from a limited region of the matrix.

### 2.12 *In vivo* PDAC model

All animal experiments were reviewed and approved by the Stony Brook University Institutional Animal Care and Use Committee (IACUC) and conducted in accordance with institutional and national guidelines for the care and use of laboratory animals. Humane endpoints included body weight loss greater than 15%, impaired mobility, severe distress, and other IACUC-approved criteria.

To establish a metastatic tumor model, 8-week-old NSG mice were purchased from The Jackson Laboratory (JAX: 005557) and were inoculated with 2 x 106 Hs 766T-Luc cells suspended in 0.2 mL PBS via intravenous (IV) tail vein injection. Following confirmation of tumor-establishment by bioluminescence IVIS imaging, mice were randomized into vehicle-control and MTX-5-FU-Gem-miR-15a treatment group. Animals were randomized (n = 6 per group; 3 male + 3 female per group) to receive either 80 µg of PEI-MAX in 5% D-glucose (vehicle control) or 80 µg of MTX-5-FU-Gem-miR-15a (4 mg/kg of body weight) formulated with 80 µg of PEI-MAX administered by tail vein injection on alternate days beginning on day 3, for a total of eight doses through day 17.

Tumor growth was monitored by luciferase expression using the IVIS Imaging System. Mice were injected with 100 µL of IVISBrite D-Luciferin, and luciferase signal was measured 10 minutes later. Tumor burden was expressed as total flux at each measurement divided by total flux at the initial post-randomization measurement. Imaging continued to day 37, twenty days after the final dose. Body weights were measured to assess treatment-associated toxicity.

### 2.13 Quantification and statistical analysis

Experiments were performed in biological triplicates unless otherwise specified; the number of independent biological replicates is stated in each figure legend. Data were analyzed using GraphPad Prism 9 and are presented as mean ± SD.

For viability, cell-cycle, and apoptosis experiments in which treatment arm was compared against the negative control within a single cell line, comparisons were made by two-tailed Student T test. For *in vivo* studies, bioluminescence measurements were analyzed using two-way ANOVA with mixed-effects modeling with treatment and time as fixed effects and repeated measurements within individual animals accounted for in the model. Statistical significance was defined as p < 0.05, indicated as p < 0.05 (*), p < 0.01 (**), p < 0.001 (***), and p < 0.0001 (****).

## 3. Results

### 3.1 MTX-5-FU-Gem-miR-15a is cytotoxic at low nanomolar concentrations across PDAC cell lines without a delivery vehicle

MTX-5-FU-Gem-miR-15a was generated by substituting all uridine and cytidine residues within the hsa-miR-15a guide strand with 5-fluorouracil and gemcitabine, and conjugating methotrexate to the 5′ end of the passenger strand (Fig. 1B). The substitutions preserve Watson-Crick base pairing, so that a single duplex carries the miR-15a guide sequence together with three antimetabolite activities.

Cytotoxicity was first evaluated under vehicle-free conditions across five PDAC cell lines representing different genetic backgrounds and previously described classical and basal-subtypes. MTX-5-FU-Gem-miR-15a reduced cell viability at low nanomolar concentrations in all five cell lines, including basal-like-associated MIA PaCa-2 and PANC-1 cells and the non-mesenchymal/classical-associated AsPC-1 and BxPC-3 models, with IC₅₀ values ranging from 0.6 ± 0.4 to 4.8 ± 0.5 nM (Fig. 1C–G; Table 1). These findings indicate that M15a retains cytotoxic activity across PDAC cell lines with distinct subtype-associated phenotypes. Importantly, no transfection reagent was required to achieve this activity under the tested conditions.

**Table 1.** IC_50_ values of MTX-5-FU-Gem-miR-15a and free drug controls across parental, gemcitabine-resistant, and daraxonrasib-resistant PDAC models.

| **Cell line** | **MTX-5-FU-Gem-miR-15a<br>IC<sub>50</sub>(nM)** | **MTX+5-FU+Gem<br>IC<sub>50</sub>(nM)** | **Fold enhancement** | **Resistance index** |
|---|---|---|---|---|
| **AsPC-1** | **4.8 ± 0.5** | **749 ± 142.7** | **~160** |  |
| **PANC-1** | **0.8 ± 0.2** | **1,091 ± 351** | **~1300** |  |
| **BxPC-3** | **0.6 ± 0.4** | **309 ± 71.8** | **~500** |  |
| **Hs 766T** | **3.9 ± 1.0** | **2,241 ± 710** | **~580** |  |
| **MIA PaCa-2** | **2.8 ± 0.2** | **362 ± 78.7** | **~130** |  |
| **Hs 766T/ GemR** | **8.6 ± 2.7** | **~ 27000 (Out of Range)** | **~3000** | **~30** |
| **MIA PaCa-2/ GemR** | **8.3 ± 0.5** | **2805 ± 227.5** | **~326** | **~25** |
| **MIA PaCa-2/ DaraxR** | **8.1 ± 2.2** | **~ 21000 (Out of Range)** | **~** | **~50** |

Data are presented as mean ± SD. IC<sub>50</sub>, half-maximal inhibitory concentration; GemR, gemcitabine-resistant; DaraxR, daraxonrasib-resistant; Resistance index, fold difference in gemcitabine/daraxonrasib IC<sub>50</sub> between parental and resistant cells

To separate the contribution of the guide sequence from that of the nucleoside-analog payload, cells were treated in parallel with free drug combination (MTX-5-FU-Gem) at 7:3:1 ratio to match the stoichiometry of the modified miRNA. These free-drug combinations required high-nanomolar to micromolar concentrations, approximately 130- to 1,400-fold less potent than MTX-5-FU-Gem-miR-15a (Fig. 1C–G; Table 1). This separation indicates that antimetabolite content alone does not account for the intact mimic’s potency.

### 3.2 MTX-5-FU-Gem-miR-15a disrupts cell-cycle distribution, induces apoptosis, and suppresses canonical miR-15a targets

We next asked how the construct affects cell-cycle distribution and cell death, and whether these effects could be attributed to the guide strand or to the antimetabolite payload. Cells were treated for 72 hours with MTX-5-FU-Gem-miR-15a (50 nM), unmodified miR-15a (50 nM), or a stoichiometrically matched free-drug combination delivering the same analog content (5-FU 350 nM, gemcitabine 150 nM, MTX 50 nM).

Flow cytometric analysis 72 hours after treatment revealed S-phase accumulation with a corresponding reduction in the G₂/S ratio across all five cell lines (Fig. 2A–B). The effect was most pronounced in MIA PaCa-2, where the ratio fell approximately 14-fold (p < 0.0001), and was similarly significant in Hs 766T (∼2.4-fold, p < 0.0001), AsPC-1 (∼3.3-fold, p < 0.0001), PANC-1 (∼3.6-fold, p < 0.0001), and BxPC-3 (∼5-fold, p < 0.01). The stoichiometrically matched free-drug combination (5-FU 350 nM, Gem 150 nM, MTX 50 nM) also lowered the G₂/S ratio, to a comparable extent in MIA PaCa-2 and BxPC-3 but only partially in Hs 766T, AsPC-1, and PANC-1, indicating that S-phase accumulation is driven largely by the antimetabolite payload with an additional contribution from the intact construct.

**Figure 2.**
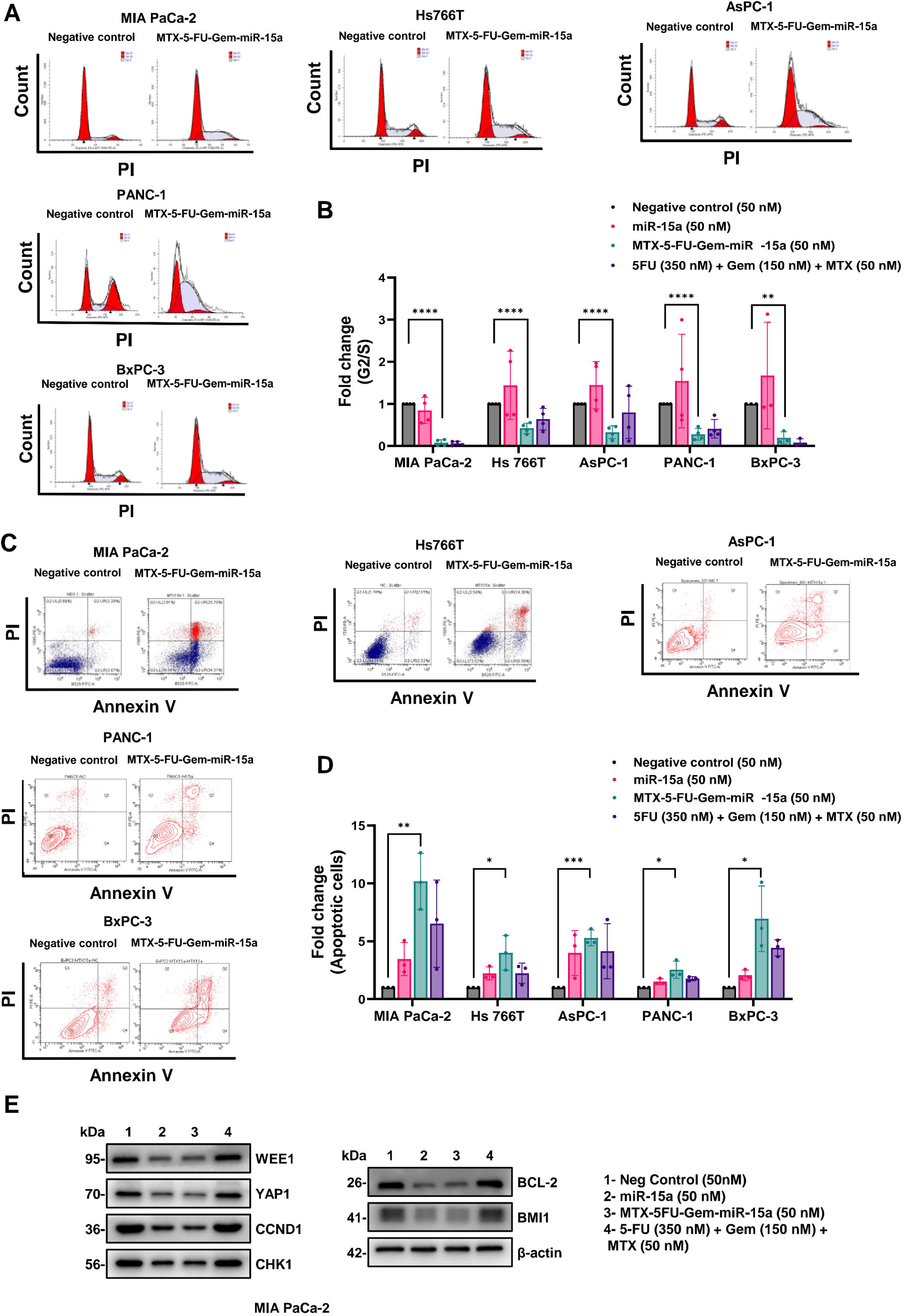
MTX-5-FU-Gem-miR-15a induces S-phase accumulation, apoptosis, and suppression of canonical miR-15a targets. (A) Representative DNA content histograms (PI staining) showing cell-cycle distribution in MIA PaCa-2, Hs 766T, AsPC-1, PANC-1, and BxPC-3 cells treated with negative control or MTX-5-FU-Gem-miR-15a. (B) Quantification of cell-cycle distribution shown as fold change in G2/S ratio relative to negative control following treatment with unmodified miR-15a (50 nM), MTX-5-FU-Gem-miR-15a (50 nM), or a stoichiometrically matched free-drug combination of 5-FU (350 nM), Gem (150 nM), and MTX (50 nM). (C) Representative Annexin V/PI flow cytometry plots showing apoptosis in the indicated PDAC cell lines following negative-control or MTX-5-FU-Gem-miR-15a treatment. (D) Quantification of total apoptotic cells (early + late apoptosis) shown as fold change relative to negative control. (E) Western blot analysis of established miR-15a target proteins in MIA PaCa-2 cells following treatment with negative control, unmodified miR-15a, MTX-5-FU-Gem-miR-15a, or the matched free-drug combination; β-actin or β-tubulin serves as a loading control. Data represent mean ± SD from n = 3 biological replicates. Statistical significance between indicated groups was determined using two-tailed Student’s t-test. ns, not significant; *p < 0.05; **p < 0.01; ***p < 0.001; ****p < 0.0001.

MTX-5-FU-Gem-miR-15a also significantly induced apoptosis (Fig. 2C–D). Annexin V/PI staining revealed increased total apoptotic populations across the panel: ∼10-fold in MIA PaCa-2 (p < 0.01), ∼3.3-fold in Hs 766T (p < 0.05), ∼5.3-fold in AsPC-1 (p < 0.001), ∼2.5-fold in PANC-1 (p < 0.05), and ∼6.9-fold in BxPC-3 (p < 0.05). In every cell line, the modified miRNA produced a greater apoptotic response than the matched free-drug combination.

To confirm that the modified mimic retained miR-15a target specificity, we examined established oncogenic miR-15a target proteins by western blot analysis. In MIA PaCa-2 cells, MTX-5-FU-Gem-miR-15a reduced the expression of CCND1, YAP1, WEE1, BCL-2, and BMI1, with a more modest effect on CHK1 (Fig. 2E). Downregulation of selected miR-15a targets was also observed in PANC-1 and AsPC-1 cells. The pattern was comparable to that produced by unmodified miR-15a delivered by transfection, whereas the matched free-drug combination left all six proteins at or above control levels, demonstrating that target suppression is directed by the guide-strand sequence rather than by the nucleoside analogs. These data indicate that extensive 5-FU and gemcitabine substitution within the guide strand preserves canonical miRNA silencing activity. Densitometric quantification is provided in Fig. S2.

Together, these findings demonstrate that MTX-5-FU-Gem-miR-15a retains canonical miR-15a-mediated target regulation while simultaneously inducing S-phase accumulation and apoptotic cell death.

### 3.3 MTX-5-FU-Gem-miR-15a retains nanomolar potency in gemcitabine-resistant PDAC cells

To assess whether MTX-5-FU-Gem-miR-15a retained activity following acquired gemcitabine resistance, resistant derivatives of MIA PaCa-2 and Hs 766T cells were generated by stepwise exposure to increasing gemcitabine concentrations. Both derivatives remained markedly less sensitive to gemcitabine than their parental counterparts after drug withdrawal, with IC₅₀ values increasing from 1.8 µM to 46.5 µM in MIA PaCa-2/GemR (∼25-fold) and from 1.9 µM to ∼55 µM in Hs 766T/GemR (∼30-fold) (Fig. S1A–B; Table 1), confirming that resistance was stable rather than a transient adaptation to ongoing selection.

Despite acquired gemcitabine resistance, MTX-5-FU-Gem-miR-15a retained low-nanomolar cytotoxic activity in both resistant derivatives, with IC₅₀ values of 8.6 ± 2.7 nM in Hs 766T/GemR and 8.3 ± 0.5 nM in MIA PaCa-2/GemR cells (Fig. 3A–B; Table 1). These values represent approximately 2.2- and 3.1-fold increases relative to the corresponding parental cells, indicating only a modest reduction in sensitivity to the construct despite pronounced gemcitabine resistance. The matched free-drug combination remained considerably less potent in both resistant models.

**Figure 3.**
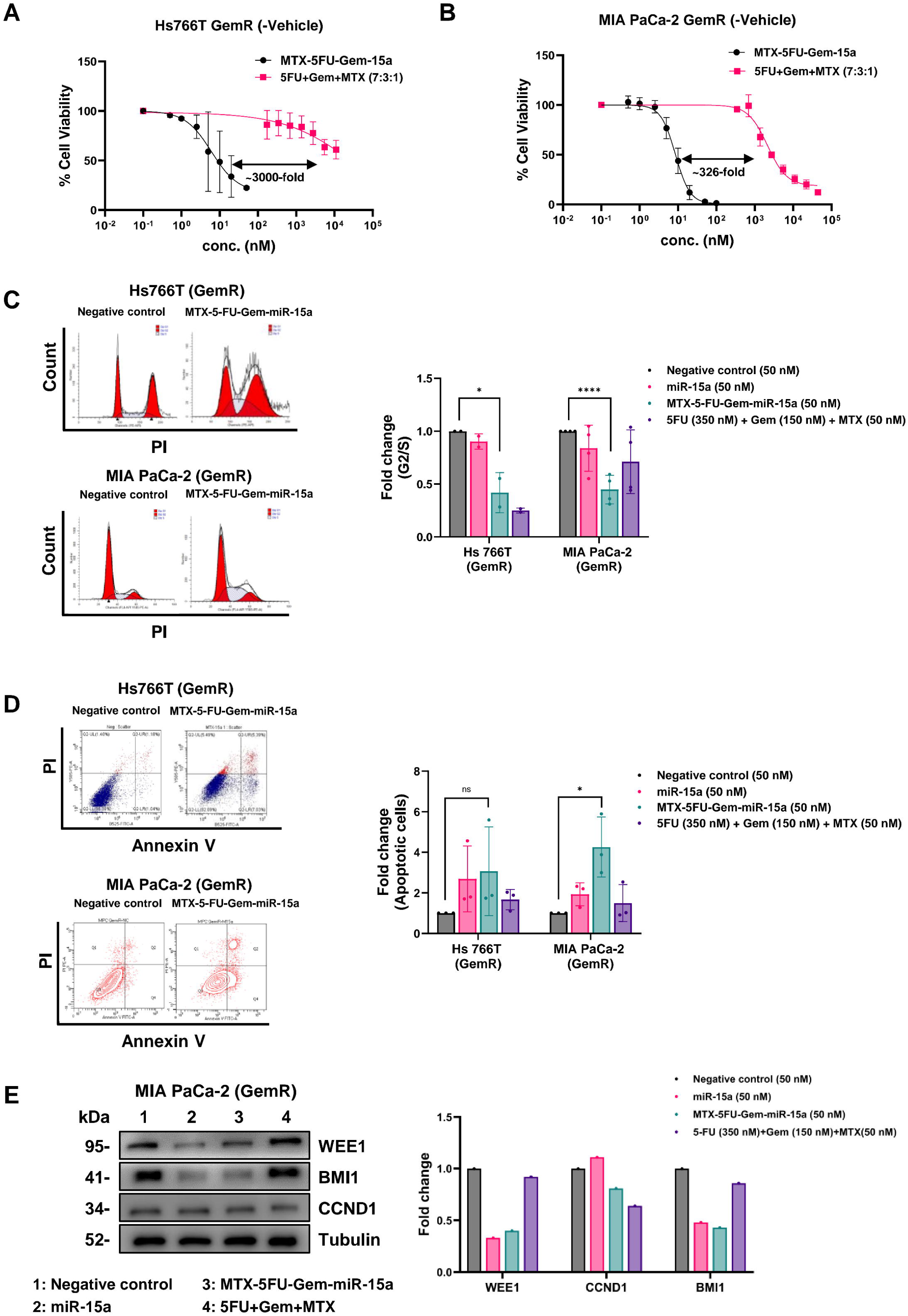
MTX-5-FU-Gem-miR-15a retains cytotoxic activity in gemcitabine-resistant pancreatic cancer cells. (A–B) Dose-response curves showing cell viability following vehicle-free treatment with MTX-5-FU-Gem-miR-15a or a stoichiometrically matched free-drug combination of 5-FU, Gem, and MTX (7:3:1) in gemcitabine-resistant Hs 766T/GemR (A) and MIA PaCa-2/GemR (B) cells. (C) Representative DNA content histograms (PI staining) and quantification of fold change in G2/S ratio following treatment with negative control, unmodified miR-15a, MTX-5-FU-Gem-miR-15a or matched free drug controls in both resistant cell lines. (D) Representative Annexin V/PI flow cytometry plots and quantification of total apoptotic cells (early + late apoptosis) shown as fold change relative to negative control. Data represent mean ± SD from n = 3 biological replicates. Statistical significance between indicated groups was determined using two-tailed Student’s t-test. ns, not significant; *p < 0.05; ****p < 0.0001.

Consistent with the effects observed in parental cells, MTX-5-FU-Gem-miR-15a treatment significantly reduced the G₂/S ratio in both resistant models, to approximately 0.4-fold in Hs 766T/GemR (p < 0.05) and 0.5-fold in MIA PaCa-2/GemR cells (p < 0.0001) (Fig. 3C). The matched free-drug combination produced a comparable reduction in Hs 766T/GemR cells but a smaller effect in MIA PaCa-2/GemR cells. The number of apoptotic cells also increased following MTX-5-FU-Gem-miR-15a treatment, approximately 4-fold in MIA PaCa-2/GemR cells (p < 0.05) and 3-fold in Hs 766T/GemR cells (p < 0.05) (Fig. 3D).

Together, these results demonstrate that MTX-5-FU-Gem-miR-15a retains potent activity following acquired gemcitabine resistance and continues to disrupt cell-cycle progression and induce apoptosis in resistant PDAC cells.

### 3.4 MTX-5-FU-Gem-miR-15a retains activity in daraxonrasib-resistant PDAC cells and synergizes with daraxonrasib

To determine whether MTX-5-FU-Gem-miR-15a also remains active against resistance arising from a mechanistically distinct therapy, a daraxonrasib-resistant derivative of MIA PaCa-2 was generated by continuous drug selection. Dose-response analysis showed that MIA PaCa-2/DaraxR viability did not fall below 50% across the daraxonrasib concentration range tested, compared with a low-nanomolar IC₅₀ in parental cells (Supplementary Fig. S1C; Table 1).

MTX-5-FU-Gem-miR-15a retained low-nanomolar cytotoxic activity in MIA PaCa-2/DaraxR cells, with an IC₅₀ of 8.1 ± 2.2 nM under vehicle-free conditions (Fig. 4A; Table 1). In contrast, the matched 5-FU, Gem, and MTX combination did not achieve 50% inhibition within the tested concentration range, showing more than 1000-fold less potency compared to the modified miRNA.

**Figure 4.**
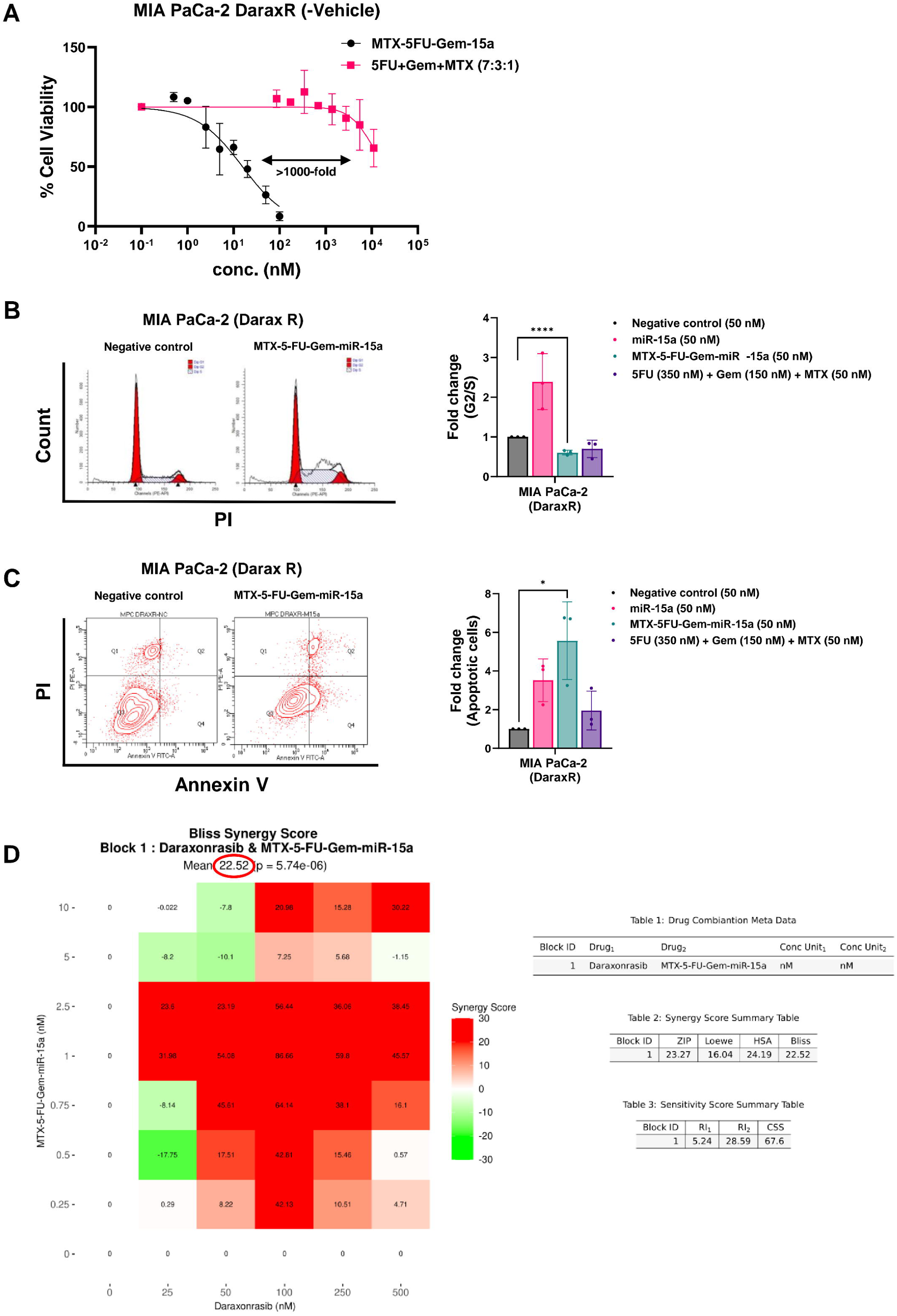
MTX-5-FU-Gem-miR-15a retains activity in daraxonrasib-resistant pancreatic cancer cells and exhibits synergy with daraxonrasib. (A) Dose-response curves showing cell viability following vehicle-free treatment with MTX-5-FU-Gem-miR-15a or a stoichiometrically matched free-drug combination of 5-FU, Gem, and MTX (7:3:1) in daraxonrasib-resistant MIA PaCa-2/DaraxR cells. (B) Representative DNA content histograms (PI staining) and quantification of fold change in G2/S ratio following treatment with negative control, unmodified miR-15a, MTX-5-FU-Gem-miR-15a, or matched free drug controls. (C) Representative Annexin V/PI flow cytometry plots and quantification of total apoptotic cells (early + late apoptosis) shown as fold change relative to negative control. (D) (D) Bliss synergy analysis of MTX-5-FU-Gem-miR-15a and daraxonrasib across the indicated concentration ranges, showing a mean Bliss synergy score of 22.52 (p = 5.74 × 10⁻). Red and green indicate synergistic and antagonistic interactions, respectively. Cell-cycle and apoptosis analyses were performed 72 hours after treatment. Data represent mean ± SD from n = 3 biological replicates. Statistical significance between indicated groups was determined using two-tailed Student’s t test. *p < 0.05; ****p < 0.0001.

MTX-5-FU-Gem-miR-15a also significantly reduced the G₂/S ratio to approximately 0.6-fold of control (p < 0.0001) (Fig. 4B). Annexin V/PI analysis showed an approximately 5.5-fold increase in apoptotic cells following MTX-5-FU-Gem-miR-15a treatment (p < 0.05), whereas the matched free-drug combination produced an approximately 2-fold increase (Fig. 4C).

Given the retained activity of MTX-5-FU-Gem-miR-15a in daraxonrasib-resistant cells, we next evaluated its interaction with daraxonrasib using a dose-response combination matrix. Bliss independence analysis revealed a strong synergistic interaction, with a mean Bliss synergy score of 22.52 (p = 5.74 × 10⁻□), and the highest synergy score of 86.66 observed at 1 nM MTX-5-FU-Gem-miR-15a and 100 nM daraxonrasib (Fig. 4D). The combination sensitivity score (CSS), which measures overall drug-combination efficacy based on dose-response inhibition, was 67.6, indicating substantial cytotoxic activity across the evaluated concentration range. Together, these findings show that MTX-5-FU-Gem-miR-15a remains active in a RAS inhibitor-resistant PDAC model and can synergize with daraxonrasib.

### 3.5 MTX-5-FU-Gem-miR-15a induces replication stress, DNA damage, and checkpoint disruption in parental and gemcitabine-resistant PDAC cells

To characterize the molecular response to MTX-5-FU-Gem-miR-15a (M15a), RNA sequencing was performed in parental (PR) and gemcitabine-resistant (GemR) MIA PaCa-2 cells treated with negative control or M15a. Mass-spectrometry-based proteomic profiling quantified approximately 5,000 proteins and provided an orthogonal approach to evaluate treatment-associated changes at the protein level. Principal component analysis clearly separated the four experimental conditions, with biological replicates clustering closely within each condition, indicating reproducible transcriptional responses. Following treatment, PR and GemR transcriptional profiles shifted closer together, suggesting partial convergence of the two cellular states (Fig. S3A).

KEGG gene set enrichment analysis revealed positive enrichment of DNA replication, Fanconi anemia, mismatch repair, homologous recombination, and cell-cycle pathways following M15a treatment in both parental and GemR cells (Fig. 5A–B). In contrast, pathways associated with oncogenic signaling and cellular metabolism, including Hippo, PI3K-Akt, mTOR, Notch, and central carbon metabolism in cancer, were negatively enriched. Proteomic analysis of parental cells showed a concordant increase in the relative abundance of proteins associated with DNA replication, mismatch repair, homologous recombination, base-excision repair, nucleotide-excision repair, and cell-cycle regulation. Apoptosis- and necroptosis-associated protein sets were also positively enriched, whereas TGF-β and Wnt signaling were negatively enriched (Fig. 5C). Together, these findings indicate a coordinated increase in the expression of replication- and repair-associated programs alongside reduced representation of selected oncogenic signaling pathways.

**Figure 5.**
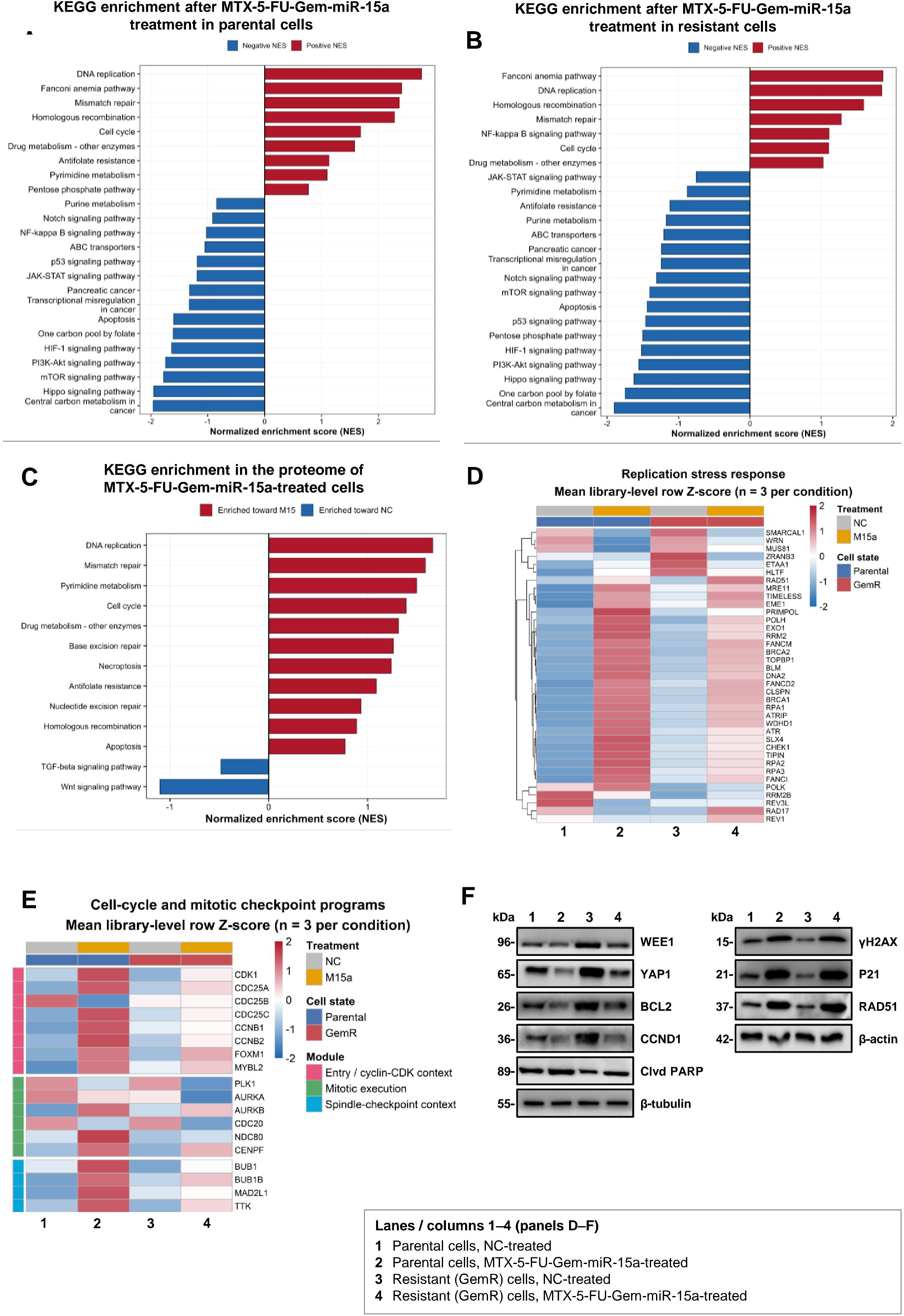
MTX-5-FU-Gem-miR-15a induces DNA damage-associated programs with checkpoint protein suppression in parental and gemcitabine-resistant PDAC cells. (A–B) KEGG pathway enrichment analyses of RNA-seq data from parental (A) and gemcitabine-resistant (B) MIA PaCa-2 cells following MTX-5-FU-Gem-miR-15a treatment, showing selected cancer-related pathways. Positive and negative normalized enrichment scores (NES) indicate enrichment toward treated and control conditions, respectively. (C) KEGG pathway enrichment analysis of quantitative proteomic data from parental MIA PaCa-2 cells showing treatment-associated changes in DNA replication, DNA repair, cell-cycle, apoptosis-associated, and signaling pathways. (D) Heatmap showing expression of genes involved in replication stress sensing, fork protection, and DNA repair in parental and GemR cells following treatment. (E) Heatmap showing expression of cell-cycle regulators, mitotic execution genes, and spindle-checkpoint-associated genes across the same conditions. Heatmaps represent mean row-wise Z-scores of normalized RNA expression from n = 3 biological replicates per condition. (F) Representative western blots showing WEE1, YAP1, BCL-2, CCND1, cleaved PARP, γH2AX, p21, and RAD51 expression in parental and GemR MIA PaCa-2 cells following treatment; β-tubulin and β-actin serve as loading controls. Lanes 1–4 represent parental negative control, parental MTX-5-FU-Gem-miR-15a, GemR negative control, and GemR MTX-5-FU-Gem-miR-15a, respectively. RNA-seq analyses included n = 3 biological replicates per condition, and quantitative proteomic analyses included n = 2 samples per treatment group.

Consistent with these pathway-level changes, curated signature scoring showed coordinated increases in DNA-damage response, replication stress, E2F/replication, and mitotic/spindle-assembly checkpoint programs following treatment (Fig. S3B). In parental cells, individual GSEA confirmed enrichment of ATR-associated replication stress (NES = 2.82, FDR = 2.2 × 10⁻¹□), DNA replication (NES = 2.66, FDR = 9.2 × 10⁻¹²), E2F targets (NES = 2.44, FDR = 3.3 × 10⁻¹□), homologous recombination (NES = 1.91, FDR = 6.8 × 10⁻□), and G2/M checkpoint (NES = 1.55, FDR = 1.8 × 10⁻□), whereas MYC Targets V1 was negatively enriched (NES = −1.94, FDR = 3.4 × 10⁻□) (Fig. S3C–H). These enrichment patterns are consistent with a transcriptional response to replication-associated stress, characterized by increased expression of DNA replication, repair, and cell-cycle regulatory genes.

Heatmap analysis showed induction of replication-stress sensing, fork-protection, and repair genes, including ATR, ATRIP, RPA1/2/3, BRCA1/2, and RAD51, with generally greater induction in parental cells (Fig. 5D). Genes associated with cell-cycle progression and mitotic regulation, including CDK1, CDC25A, CDC25C, CCNB1, CCNB2, and FOXM1, were also increased following treatment (Fig. 5E). Consistent with these transcriptional changes, proteomic analysis showed increased abundance of CDK1, CCNB1, MKI67, TOP2A, NDC80, and KIF23 in treated parental cells (Fig. S4A). Phosphoproteomic profiling also showed increased LMNA pS22, MCM3 pS668/pS672, PAK2 pS141, and CFL1 pS3, reflecting treatment-associated changes in proteins involved in replication stress, mitotic progression, and cell death (Fig. S4B).

In contrast to the induction of replication and repair programs, checkpoint regulation showed a distinct pattern. Although WEE1 and CHK1 transcripts increased, western blot analysis demonstrated reduced WEE1 and CHK1 proteins in both parental and resistant cells (Fig. 5F). This divergence is consistent with compensatory transcription during replication stress alongside maintained miR-15a-mediated protein suppression. YAP1, BCL-2, and CCND1, which were elevated in untreated GemR cells relative to parental cells, were similarly reduced following treatment. Western blot analysis further confirmed treatment-associated DNA damage, with increased γH2AX in both backgrounds, particularly in GemR cells, together with increased RAD51 and p21. Cleaved PARP increased most clearly in GemR cells, whereas parental cells showed comparatively high basal cleaved PARP (Fig. 5F).

Proteomic analysis further supported the apoptotic response, with increased death-receptor and execution-associated proteins, including FADD, TNFRSF10B, PARP1, and DFFA, as well as mitochondrial death-response proteins CYCS, AIFM1, DNM1L, and OPA1 (Fig. S4C). Together with the increased Annexin V-positive populations observed in functional assays, these changes support activation of apoptotic cell death despite the absence of corresponding positive transcriptomic apoptosis enrichment. This difference further highlights the importance of protein-level responses in characterizing the effects of the modified miRNA.

Collectively, transcriptomic, proteomic, phosphoproteomic, and western blot analyses demonstrate that M15a induces replication-associated DNA damage and repair responses while simultaneously reducing checkpoint and survival proteins. The concurrent induction of replication, repair, and mitotic programs, together with reduced WEE1 and CHK1 and γH2AX accumulation, is consistent with continued cell-cycle progression in the presence of unresolved replication-associated damage.

### 3.6 MTX-5-FU-Gem-miR-15a remodels gemcitabine resistance-associated programs

Having established the molecular response shared between parental and resistant cells, we next examined whether MTX-5-FU-Gem-miR-15a altered pathways specifically associated with acquired gemcitabine resistance.

Pathway-level analysis identified substantial alterations in drug-handling and antimetabolite-response programs in untreated GemR cells (Fig. 6A). ABC transporters, antifolate resistance, drug metabolism-other enzymes, purine metabolism, and pyrimidine metabolism showed positive enrichment in the resistant state, whereas one-carbon pool by folate showed negative enrichment. Following M15a treatment, several pathways shifted in the opposite direction, including ABC transporters, antifolate resistance, and purine and pyrimidine metabolism, indicating broad remodeling of pathways associated with gemcitabine resistance.

**Figure 6.**
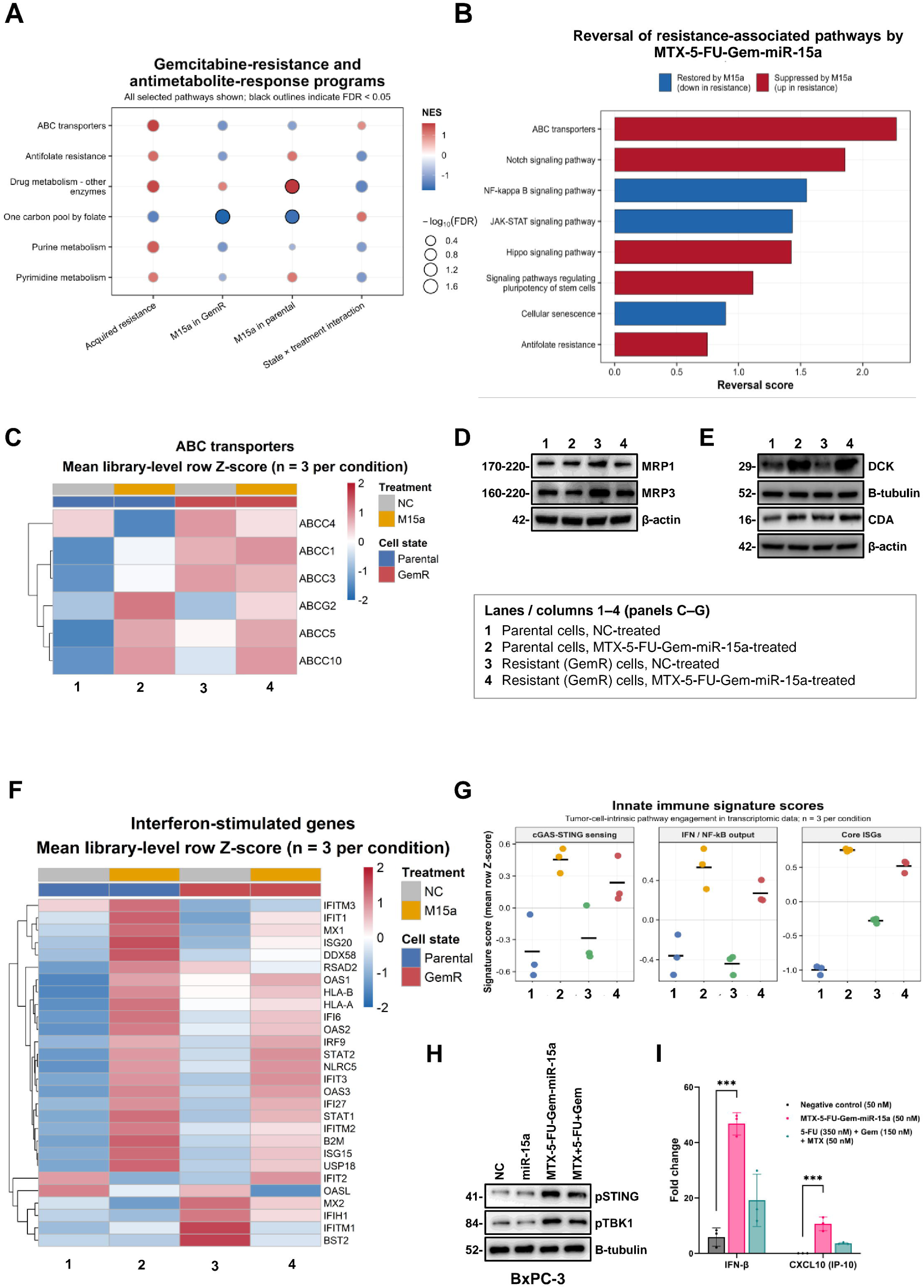
MTX-5-FU-Gem-miR-15a remodels gemcitabine resistance-associated programs and induces innate immune signaling. (A) KEGG pathway enrichment analysis of drug-handling and antimetabolite-response pathways in untreated GemR cells relative to parental controls and following MTX-5-FU-Gem-miR-15a treatment in parental and resistant cells. The state × treatment interaction is also shown. Circle color represents normalized enrichment score (NES), circle size represents −log10(FDR), and black outlines indicate FDR < 0.05. (B) Pathway-reversion analysis showing resistance-associated signaling programs opposed by MTX-5-FU-Gem-miR-15a treatment. Positive reversion scores indicate a treatment-associated shift opposite to the resistant-state direction. (C) Heatmap showing expression of selected ABC transporter genes in parental and GemR cells before and after treatment. (D) Western blot analysis of MRP1/ABCC1 and MRP3/ABCC3 in parental and GemR cells following treatment; β-actin serves as a loading control. (E) Western blot analysis of deoxycytidine kinase (dCK) and cytidine deaminase (CDA) in parental and GemR cells following treatment; α-tubulin and β-actin serve as loading controls. (F) Heatmap showing expression of interferon-stimulated and antigen-presentation genes in parental and GemR cells under control and treatment conditions. (G) Curated signature scores showing cGAS-STING sensing, IFN/NF-κB output, and core interferon-stimulated gene (ISG) programs across the four experimental conditions. Heatmaps and signature scores represent RNA-seq data from n = 3 biological replicates per condition. For panels C–G, conditions 1–4 represent parental negative control, parental MTX-5-FU-Gem-miR-15a, GemR negative control, and GemR MTX-5-FU-Gem-miR-15a, respectively. (H) Western blot analysis of phosphorylated STING (p-STING) and TBK1 (p-TBK1) in BxPC-3 cells following treatment with negative control, unmodified miR-15a, MTX-5-FU-Gem-miR-15a or matched free drug controls; α-tubulin serves as the loading control. (I) ELISA measurement of secreted IFN-β and CXCL10 in conditioned media from gemcitabine-resistant cells following MTX-5-FU-Gem-miR-15a treatment.

Among 33 curated resistance-associated genes, 14 shifted in the opposite direction to that observed during acquisition of resistance following M15a treatment (Fig. S5A). Notch-associated genes JAG1, NRARP, CCND1, and HES1 were reduced, together with matrix- and EMT-associated genes SPARC, SNAI2, and CTGF, further supporting suppression of resistance-associated signaling and cell-state programs.

Pathway-reversion analysis further identified ABC transporters, Notch, NF-κB, JAK-STAT, Hippo, pluripotency, cellular senescence, and antifolate resistance among the programs opposed by M15a treatment (Fig. 6B). Pathways elevated during resistance, including ABC transporters, Notch, Hippo, pluripotency, and antifolate resistance, were suppressed, whereas NF-κB, JAK-STAT, and cellular senescence-associated programs shifted toward restoration. GSEA showed a directional change in ABC transporter signaling, from positive enrichment during resistance (NES = 1.56, FDR = 0.060) to negative enrichment following M15a treatment (NES = −1.22, FDR = 0.406) (Fig. S5B–C). Pyrimidine metabolism showed a similar shift, from NES = 1.08 (FDR = 0.291) to NES = −1.01 (FDR = 0.424) (Fig. S5D–E). Because these individual comparisons did not reach FDR significance, the findings suggest potential remodeling rather than definitive reversal of the corresponding pathways.

We next examined whether the treatment-associated changes in ABC transporter signatures were reflected at the protein level. Transcriptomic analysis identified distinct expression patterns among individual ABC transporter genes in parental and GemR cells (Fig. 6C). Western blot analysis revealed that MRP1/ABCC1 and MRP3/ABCC3 were strongly elevated in untreated GemR cells relative to parental cells and markedly reduced following M15a treatment (Fig. 6D). These protein-level changes were greater than the corresponding transcript responses, where ABCC1 showed little change and ABCC3 was only modestly reduced. ABCC4, another transporter associated with nucleotide-metabolite efflux, was also reduced at the transcript level (Fig. S5A).

We next examined genes involved in gemcitabine transport and metabolism. DCK, which encodes the enzyme responsible for the initial phosphorylation of gemcitabine, increased following M15a treatment in GemR cells (log₂-fold change ≈ +1.15), whereas SLC29A1 expression remained largely unchanged (Fig. S5A). Consistent with these findings, western blot analysis showed reduced dCK protein abundance in untreated GemR cells relative to parental cells and increased dCK expression following M15a treatment (Fig. 6E). This restoration of dCK, with only modest CDA induction, is consistent with a shift in the balance between gemcitabine activation and inactivation, although functional resensitization was not assessed.

Together, these findings indicate that M15a remodels drug-responding and resistance-associated transcription state in GemR cells, with particularly pronounced effects on the efflux proteins MRP1/MRP3 and the gemcitabine-activating enzyme dCK.

### 3.7 MTX-5-FU-Gem-miR-15a induces an interferon-associated innate immune response in parental and gemcitabine-resistant PDAC cells

Transcriptomic analysis also revealed a pronounced interferon-associated response following M15a treatment in parental and GemR MIA PaCa-2 cells. Interferon-stimulated genes, including IFIT1, IFIT3, MX1, MX2, ISG15, OAS1, OAS2, OAS3, RSAD2, STAT1, STAT2, and IRF9, were increased following treatment, together with antigen-presentation-associated genes HLA-A, HLA-B, and B2M (Fig. 6F). Curated signature analysis showed increased expression of genes associated with cGAS-STING signaling, interferon/NF-κB responses, and core interferon-stimulated programs in both parental and resistant cells (Fig. 6G). These findings demonstrate that acquired gemcitabine resistance does not prevent induction of an interferon-associated transcriptional response to M15a.

Proteomic profiling independently supported this response, with positive enrichment of cytosolic DNA sensing, innate immune system, interferon α/β and γ signaling, IL6/JAK/STAT3, TNFα/NF-κB, and inflammatory-response pathways (Fig. S4D). STAT1, DDX3X, and TOMM70 increased, together with DNA damage-associated proteins PRKDC, XRCC5, and XRCC6 (Fig. S4E). In contrast, the RIG-I/MDA5 RNA-sensing pathway was not similarly activated at the protein level, suggesting a response more closely associated with DNA damage sensing than generalized recognition of the RNA duplex.

To further examine whether M15a treatment was associated with STING-TBK1 signaling, we evaluated phosphorylation of STING and TBK1 in BxPC-3 cells. Western blot analysis showed increased p-STING and p-TBK1 following M15a treatment (Fig. 6H), providing additional evidence of treatment-associated innate immune signaling in a separate PDAC cell-line background. To determine whether this signaling produced a functional secreted response, conditioned media were analyzed for IFN-β and CXCL10. Both were significantly increased in GemR cells following treatment (Fig. 6I). Together, these findings suggest that M15a induces tumor-cell-intrinsic innate immune signaling alongside replication-associated DNA damage, consistent with activation of the cGAS-STING-TBK1 pathway.

### 3.8 MTX-5-FU-Gem-miR-15a is active in patient-derived organoids and suppresses tumor growth *in vivo*

To evaluate MTX-5-FU-Gem-miR-15a activity in patient-derived PDAC models, three organoid lines (hF3, hF44, and hT89) were treated with increasing concentrations of M15a or gemcitabine. M15a produced a dose-dependent reduction in organoid viability, with IC₅₀ values of 2.76, 2.69, and 3.97 nM in hF3, hF44, and hT89, respectively (Fig. 7C–E). In comparison, gemcitabine exhibited IC₅₀ values of approximately 420, 380, and 470 nM in the corresponding organoid lines, indicating that MTX-5-FU-Gem-miR-15a was approximately 120- to 150-fold more potent than gemcitabine. M15a treatment resulted in near-complete loss of viability in two organoid lines, whereas the third retained residual viability following both treatments. In this model, gemcitabine reached a viability plateau of approximately 15%, while M15a continued to reduce viability across the tested concentration range. Together, these findings demonstrate that MTX-5-FU-Gem-miR-15a exhibits greater cytotoxic potency than gemcitabine in patient-derived PDAC organoids.

**Figure 7.**
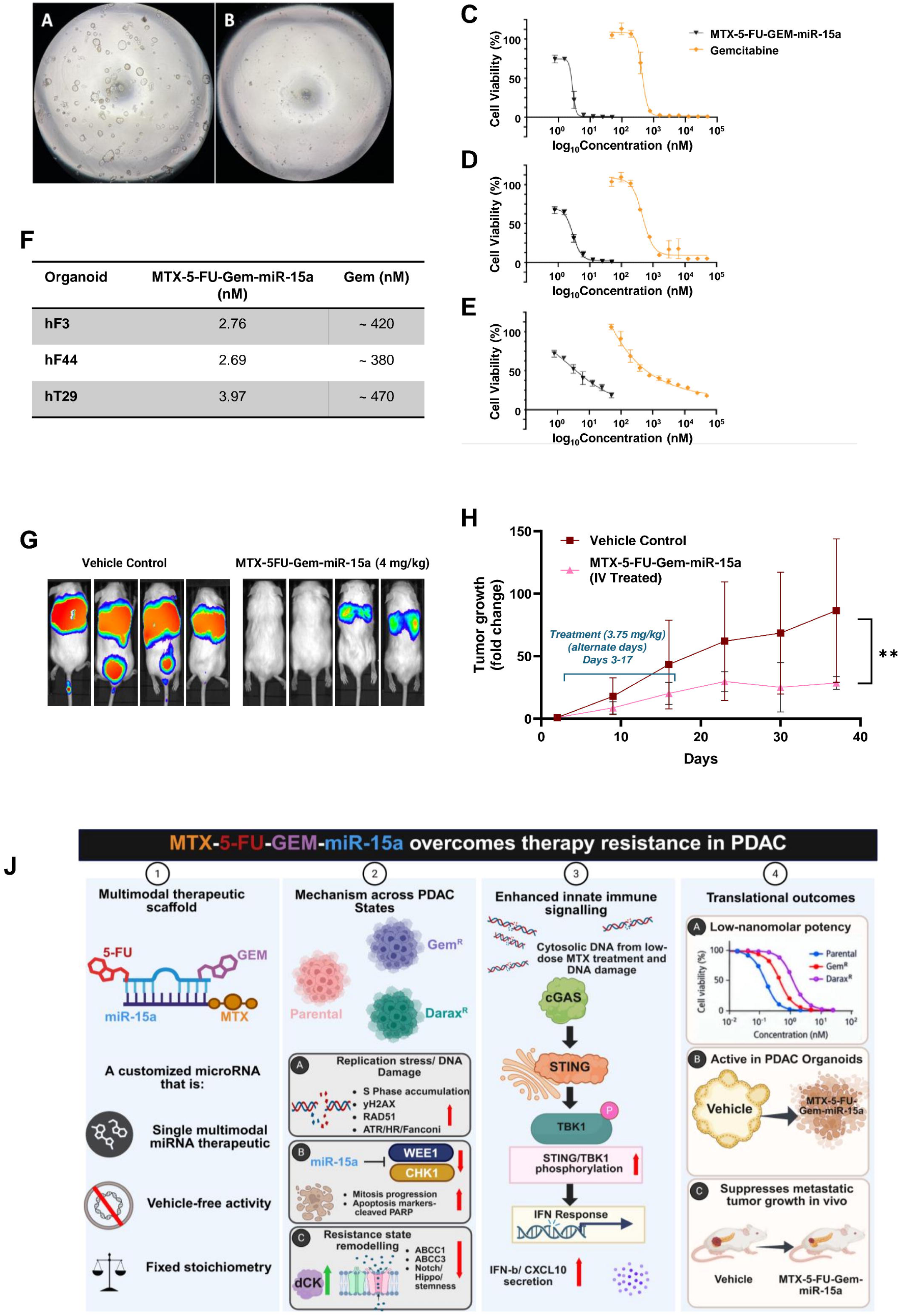
MTX-5-FU-Gem-miR-15a exhibits potent activity in patient-derived pancreatic cancer organoids and suppresses tumor growth in vivo. (A–B) Representative brightfield images of patient-derived PDAC organoids under control (A) and MTX-5-FU-Gem-miR-15a-treated (B) conditions, showing treatment-associated loss of organoid structural integrity. (C–E) Dose-response curves showing cell viability following treatment with MTX-5-FU-Gem-miR-15a or gemcitabine in three patient-derived PDAC organoid lines. (F) Summary of half-maximal inhibitory concentration (IC₅₀) values for MTX-5-FU-Gem-miR-15a and gemcitabine across the three organoid lines. (G) Representative IVIS bioluminescence images of mice bearing intravenously established Hs 766T-Luc tumors following treatment with vehicle or MTX-5-FU-Gem-miR-15a (4 mg/kg). (H) Longitudinal quantification of tumor burden shown as fold change in total bioluminescent flux relative to the initial post-randomization measurement. Dashed lines indicate the treatment schedule, with intravenous administration on alternate days from day 3 through day 17. Data are presented as mean ± SD. Tumor growth was analyzed using two-way ANOVA with mixed-effects modeling to account for repeated measurements. **p < 0.01. (J) Proposed mechanistic model of MTX-5-FU-Gem-miR-15a activity, illustrating suppression of miR-15a target proteins, remodeling of resistance-associated pathways and gemcitabine metabolism, and induction of interferon-associated innate immune signaling alongside antitumor activity in organoids and the experimental metastatic model.

To assess antitumor activity *in vivo*, 8-week-old NSG mice were inoculated with luciferase-expressing Hs 766T cells by intravenous tail-vein injection to establish a metastatic PDAC model. Following confirmation of tumor establishment by bioluminescent imaging, animals were randomized to receive vehicle or MTX-5-FU-Gem-miR-15a at 4 mg/kg by tail-vein injection every other day for a total of eight doses (Fig. 7G). Tumor burden was monitored longitudinally by IVIS imaging and quantified as fold change in total bioluminescent flux relative to the initial post-randomization measurement. MTX-5-FU-Gem-miR-15a treatment significantly suppressed tumor growth compared with vehicle controls throughout the observation period (two-way ANOVA, p < 0.01) (Fig. 7H). Separation between the treatment groups became evident during dosing and persisted for about 20 days after the final dose, indicating a sustained antitumor effect after treatment cessation. By day 37, tumor burden in vehicle-treated animals had increased approximately 85-fold from baseline, compared with approximately 33-fold in MTX-5-FU-Gem-miR-15a-treated animals, representing an approximately 60% reduction in relative tumor expansion. Endpoint IVIS imaging similarly showed substantially lower metastatic bioluminescent signal in treated animals compared with vehicle controls (Fig. 7G). Together, these findings demonstrate that the low-nanomolar activity of MTX-5-FU-Gem-miR-15a extends to patient-derived PDAC organoids and translates into sustained suppression of metastatic tumor progression *in vivo*.

## 4. Discussion

Therapeutic resistance remains a major limitation in PDAC, affecting both chemotherapy and emerging RAS(ON) targeted therapies. Gemcitabine resistance involves altered drug activation, nucleotide metabolism, efflux, and survival-associated signaling, whereas acquired resistance to RAS(ON) inhibitors frequently involves reactivation of RAS signaling [6, 7]. Despite their distinct origins, these states may share dependencies on cell-cycle regulation and DNA-damage responses. In this study, we developed MTX-5-FU-Gem-miR-15a to combine antimetabolite-induced replication stress with miR-15a-mediated suppression of oncogenic and checkpoint proteins. The study’s principal finding is that a single construct retains potent activity across two resistance states arising from mechanistically distinct therapeutic pressures, supporting a strategy that targets multiple resistance-associated vulnerabilities simultaneously. Its activity across PDAC cell lines with basal-like- and classical-associated phenotypes further suggests that sensitivity is not restricted to a single molecular phenotype, although subtype-specific differences were not directly evaluated [38, 46].

Previous work in our laboratory established miR-15a as a tumor-suppressive miRNA in PDAC, restoring miR-15a suppresses WEE1, CHK1, YAP1, and BMI1 and increases sensitivity to gemcitabine [22]. This provided the basis for developing chemically modified miR-15a with improved therapeutic activity. Previous 5-FU- and Gem-modified miR-15a constructs retained sequence-specific target regulation while improving potency and vehicle-free activity, and the present construct represents a further extension of this platform by combining 5-FU, Gem, and MTX within a single miRNA scaffold [22]. Consistent with this design, MTX-5-FU-Gem-miR-15a remained active without a transfection reagent, whereas unmodified miR-15a was inactive under vehicle-free conditions. Stoichiometrically matched free 5-FU, Gem, and MTX were substantially less potent than MTX-5-FU-Gem-miR-15a (Table 1), while the intact MTX-5-FU-Gem-miR-15a retained suppression of established miR-15a targets including WEE1, CHK1, YAP1, CCND1, BCL-2, and BMI1 (Fig. 2E). Because the antimetabolites are incorporated within the duplex rather than co-administered with it, every cell that takes up the construct receives the miR-15a guide sequence and all three antimetabolite components at a fixed stoichiometry, potentially reducing the pharmacologic separation and asynchronous exposure that occur when the same agents are given as a mixture. These findings indicate that antimetabolite exposure alone cannot explain the construct’s activity; instead, its potency likely involves combined contributions of chemical modification, intracellular delivery, and sequence-specific miR-15a activity.

Mechanistically, MTX-5-FU-Gem-miR-15a couples replication stress with suppression of the checkpoint response required to tolerate it. Gemcitabine interferes with DNA synthesis by terminating DNA chains and inhibiting ribonucleotide reductase, while 5-FU limits thymidylate synthesis and MTX disrupts folate-dependent nucleotide metabolism [30–33]. The matched antimetabolite combination induced S-phase accumulation, indicating that nucleotide disruption contributes substantially to the cell-cycle phenotype. Treatment with MTX-5-FU-Gem-miR-15a was accompanied by increased expression of genes associated with ATR-associated replication stress, Fanconi anemia, homologous recombination, and cell-cycle programs, consistent with the proteomic enrichment of DNA replication and repair pathways and accumulation of γH2AX. These transcriptional changes likely reflect a cellular response to replication-associated damage rather than increased functional DNA repair capacity. At the same time, WEE1 and CHK1 proteins decreased despite increased WEE1 and CHK1 transcripts (Fig. 5F). Because both proteins are previously validated miR-15a targets in PDAC, this divergence is consistent with compensatory transcription during replication stress occurring alongside post-transcriptional suppression [14, 22]. Reduced checkpoint protein abundance, together with increased mitotic and spindle-associated programs, suggests that cells may progress toward mitosis without adequately resolving replication-associated damage. This provides a mechanistic explanation for the observed apoptosis and is consistent with a mitotic-catastrophe-like response, although direct visualization of aberrant mitosis is needed for confirmation. Unlike direct CHK1 or WEE1 inhibition, which has faced toxicity and efficacy limitations in clinical development, the construct generates replication stress and reduces checkpoint protein expression in the same treated cells [19, 20, 47].

The multi-omic data also provide further insight into the response of resistant cells. Gemcitabine-resistant cells had higher baseline expression of replication-stress and repair programs but showed a smaller treatment-induced transcriptional response than parental cells despite retaining sensitivity. This suggests that pathway induction does not necessarily reflect the magnitude of the cytotoxic response. Furthermore, the apoptosis-associated response was more evident in proteomic and functional measurements than in transcript-level pathway enrichment. We observed similar transcript-protein differences for WEE1, CHK1, and the efflux transporters. These findings are consistent with the post-transcriptional activity of miR-15a and emphasize the importance of interpreting RNA-seq alongside protein and functional data.

In GemR cells, dCK, the enzyme responsible for the initial phosphorylation of gemcitabine, was markedly reduced, consistent with previous studies linking loss of dCK to acquired gemcitabine resistance [8, 12]. Following MTX-5-FU-Gem-miR-15a treatment, DCK expression increased, and dCK protein was restored, whereas CDA, which contributes to gemcitabine inactivation, increased only modestly (Fig. 6E) [48]. Thus, MTX-5-FU-Gem-miR-15a restored a key component of gemcitabine metabolism that was diminished during acquired resistance. Whether increased dCK abundance translates into enhanced activation of free gemcitabine and functional resensitization remains to be established. Importantly, MTX-5-FU-Gem-miR-15a itself retained low-nanomolar potency in GemR cells despite their reduced baseline dCK expression. Incorporation of Gem within an oligonucleotide may alter its cellular uptake and intracellular processing, but it is not yet known whether the incorporated residues bypass conventional dCK-mediated activation.

Drug-efflux proteins were also markedly altered by treatment. MRP1/ABCC1 and MRP3/ABCC3 were markedly increased in untreated GemR cells and strongly reduced following treatment (Fig. 6D). Increased expression of ABC transporters, including MRP1 and MRP3, has been associated with chemoresistance in PDAC, with direct evidence implicating MRP1 in gemcitabine resistance [49–51]. Thus, the downregulation of MRP1 and MRP3 by MTX-5-FU-Gem-miR-15a may contribute to partial resensitization of resistant cells to gemcitabine by attenuating resistance-associated drug-efflux pathways. The decrease in transporter proteins was more pronounced than the corresponding transcript changes, suggesting additional post-transcriptional or indirect regulation. Together with increased dCK expression, these changes suggest that M15a may partially restore gemcitabine sensitivity by targeting both drug-efflux-associated pathways and gemcitabine activation. However, functional resensitization to free gemcitabine remains to be established. Increased RRM1, RRM2, and TYMS expression following treatment may reflect a compensatory response to antimetabolite-induced nucleotide stress rather than persistence of the original resistant state [7, 52].

M15a also induced transcriptional changes opposing Notch, Hippo, pluripotency, and nucleotide-metabolism programs associated with gemcitabine resistance. Decreased expression of JAG1, NRARP, HES1, and CCND1, together with SPARC, SNAI2, and CTGF, is consistent with attenuation of Notch-associated, EMT, and stem-like programs. These pathways have been implicated in aggressive PDAC and gemcitabine resistance, and previous studies have reported enhanced gemcitabine activity following Notch inhibition. Notably, the reduction of HES1 and NRARP, both Notch-responsive genes, supports decreased pathway output rather than changes in receptor expression alone [13, 26, 53]. The treatment response therefore extends beyond drug metabolism to programs that help maintain the resistant cellular state.

The retained activity of MTX-5-FU-Gem-miR-15a in daraxonrasib-resistant MIA PaCa-2 cells is particularly relevant as RAS(ON) inhibition advances in PDAC. MIA PaCa-2 cells harbor a *KRAS* G12C mutation, providing a model to investigate acquired resistance to daraxonrasib in this genetic background. Acquired resistance to daraxonrasib has been associated with mutant *KRAS* amplification and alterations involving RTK, MAPK, and PI3K signaling, while some resistant models exhibit increased dependence on ATR/CHK1/WEE1 signaling [6]. Despite their reduced sensitivity to daraxonrasib, resistant MIA PaCa-2 cells remained sensitive to M15a, and the combination was synergistic, with a mean Bliss score of 22.52 (Fig. 4D). By inducing replication stress while reducing WEE1 and CHK1 proteins, M15a may exploit checkpoint dependencies that persist or emerge during RAS-pathway reactivation. Its effects on PI3K-Akt, mTOR, Hippo, and Notch-associated pathways provide an additional rationale for activity across different mechanisms of resistance [6, 14, 27]. Although the mechanism of acquired resistance in our DaraxR model remains undefined, these findings support further investigation of M15a in combination with daraxonrasib. Importantly, additional daraxonrasib-resistant models harboring other common *KRAS* mutations, particularly G12D and G12V, will be needed to determine whether this activity extends across different *KRAS*-mutant backgrounds.

In addition to its effects on cell survival, M15a induced a tumor-cell-intrinsic interferon response. Increased interferon-stimulated and antigen-presentation gene expression, phosphorylation of STING and TBK1, and secretion of IFN-β and CXCL10 support activation of innate immune signaling following treatment (Fig. 6H–I). Replication-associated DNA damage may contribute to this response, although direct dependence on cGAS-STING remains to be established. Low-dose MTX has been reported to activate cGAS-STING signaling in tumor cells, suggesting a possible contribution from the incorporated MTX moiety [34, 54]. Low-dose 5-FU and Gem have also been reported to affect myeloid-derived suppressor cells and tumor infiltrating regulatory T cells (TI-Tregs), respectively, raising additional questions about immune modulation [55, 56]. Whether the interferon response or the incorporated antimetabolites affect these populations will require immunocompetent models.

Activity in patient-derived organoids and an experimental metastatic model supports further therapeutic investigation. Low-nanomolar cytotoxicity in three organoid lines and suppression of metastatic tumor growth after systemic administration extend these findings beyond monolayer culture (Fig. 7H). The difference between treated and vehicle groups persisted for about 20 days after dosing ended, suggesting sustained suppression of tumor growth. However, systemic treatment used PEI-MAX formulation, and the immunodeficient model cannot establish carrier-independent delivery or antitumor immunity.

Three principal areas require further investigation. Gemcitabine rechallenge, dCK activity assays, and intracellular metabolite measurements are needed to determine whether restored dCK increases sensitivity to free gemcitabine and how the incorporated Gem residues are processed. Matched free-drug controls could help determine whether dCK restoration is specific to the modified miRNA or a secondary consequence of altered cell-cycle distribution. Direct imaging is required to establish mitotic catastrophe. The contribution of MTX to uptake, including possible FRα dependence, should be evaluated alongside pharmacokinetics, biodistribution, and penetration through PDAC stroma. Finally, studies in immunocompetent and drug-resistant tumor models are needed to define effects on immune recruitment, TI-Tregs, MDSCs, and the durability of therapeutic activity.

In summary, MTX-5-FU-Gem-miR-15a offers a unique approach to treat therapy-resistant PDAC by combining antimetabolite-induced replication stress with miR-15a-mediated suppression of cell cycle checkpoints, oncogenic drivers, and survival proteins. This enhanced therapeutic activity was reflected in low-nanomolar cytotoxicity and apoptosis; replication stress induction alongside checkpoint disruption; remodeling of gemcitabine resistance-associated metabolic, efflux, and cell-state programs; activation of tumor-cell-intrinsic innate immune signaling; and sustained activity in patient-derived organoids and in vivo. The retention of sensitivity across gemcitabine- and daraxonrasib-resistant PDAC model systems converges on impacting complicated adaptive survival and DNA-damage response mechanisms. By combining induction of replication stress with simultaneous suppression of the checkpoint machinery required to survive that stress, MTX-5-FU-Gem-miR-15a provides a single therapeutic platform capable of engaging vulnerabilities shared across mechanistically distinct resistance states. This study establishes a solid foundation for further evaluation of MTX-5-FU-Gem-miR-15a as a novel therapeutic agent to overcome resistance mechanisms in PDAC.

## CRediT authorship contribution statement

A.P. and J.J. conceptualized the study. A.P., A.O., J.G.Y., H.B., L.C., and I.O. developed methodology. A.P., A.O., and J.G.Y. performed experiments. A.P. performed formal analysis, including RNA-sequencing and bioinformatic analyses. H.B., L.C., and I.O. contributed chemical synthesis. A.P. wrote the original draft. A.P., A.O., and J.J. reviewed and edited the manuscript. J.J. supervised the study, acquired funding, and provided resources. All authors read and approved the final manuscript.

## Ethics approval

All animal experiments were reviewed and approved by the Stony Brook University Institutional Animal Care and Use Committee (IACUC; protocol 293153) and were conducted in accordance with institutional policies. Humane endpoints included body weight loss greater than 15%, impaired mobility, severe distress, or other IACUC-approved endpoint criteria, and euthanasia was performed in accordance with AVMA guidelines. Patient-derived pancreatic ductal adenocarcinoma organoids (hF3, hF44, and hT89) were provided by the Stony Brook Medicine Biobank (Stony Brook Medicine Biobank, Renaissance School of Medicine, Stony Brook, USA).

## Data availability

RNA-sequencing data generated in this study will be deposited in the Gene Expression Omnibus and will be publicly available as of the date of publication. All other data supporting the findings of this study are included in the article and supplementary information files or are available from the corresponding author upon reasonable request. MTX-5-FU-Gem-miR-15a and related reagents generated in this study will be made available by the lead contact upon reasonable request and may require a completed material transfer agreement. Code used for RNA-sequencing and downstream analyses will be made publicly available upon publication or provided upon reasonable request.

## Acknowledgements

We thank members of the Ju laboratory and the Department of Pathology for helpful discussions and technical assistance. Mass spectrometry analyses were performed by the Stony Brook University Proteomics Center.

## Declaration of generative AI and AI-assisted technologies in the writing process

During preparation of this manuscript, the authors used ChatGPT, developed by OpenAI, for language editing and manuscript organization. All scientific content, data interpretation, and conclusions were reviewed, edited, and approved by the authors, who take full responsibility for the final manuscript.

## Supplementary information

**Additional file 1: Document_S1.pdf**

Supplementary Figures S1–S5.

**Additional file 2: Table_S2.csv**

Differential gene expression of parental and Gemcitabine-resistant MIA PaCa-2 cells following MTX-5-FU-Gem-miR-15a treatment, related to Fig. 5 and Fig. 6.

