## Supplemental Data 1 for "Developing a Multimodal miR-15a Mimic to Overcome Gemcitabine and RAS(ON) Inhibitor Daraxonrasib Resistance in Pancreatic Ductal Adenocarcinoma"

### Slide 1
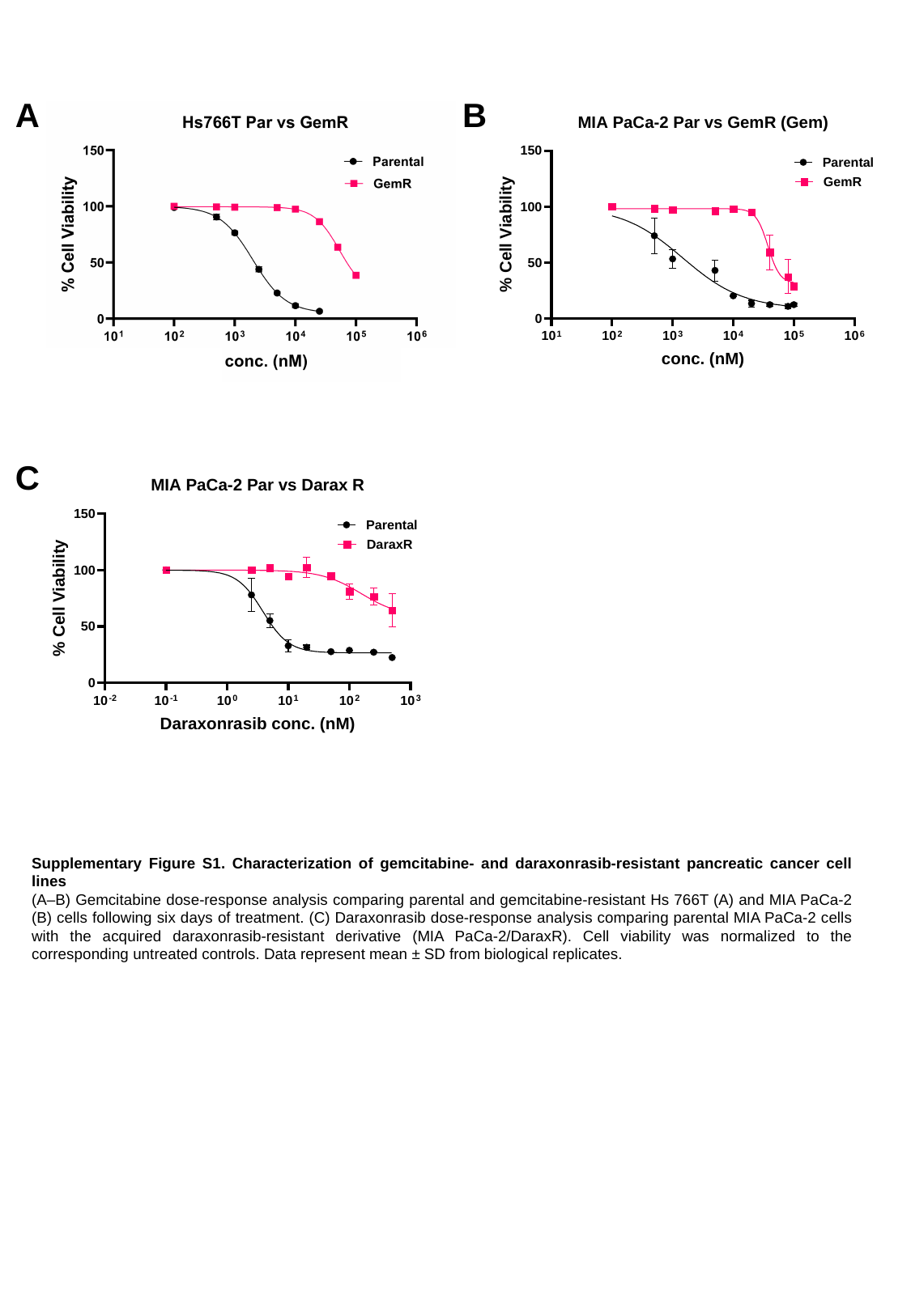

A
B
C
Supplementary Figure S1. Characterization of gemcitabine- and daraxonrasib-resistant pancreatic cancer cell lines
(A–B) Gemcitabine dose-response analysis comparing parental and gemcitabine-resistant Hs 766T (A) and MIA PaCa-2 (B) cells following six days of treatment. (C) Daraxonrasib dose-response analysis comparing parental MIA PaCa-2 cells with the acquired daraxonrasib-resistant derivative (MIA PaCa-2/DaraxR). Cell viability was normalized to the corresponding untreated controls. Data represent mean ± SD from biological replicates.

### Slide 2
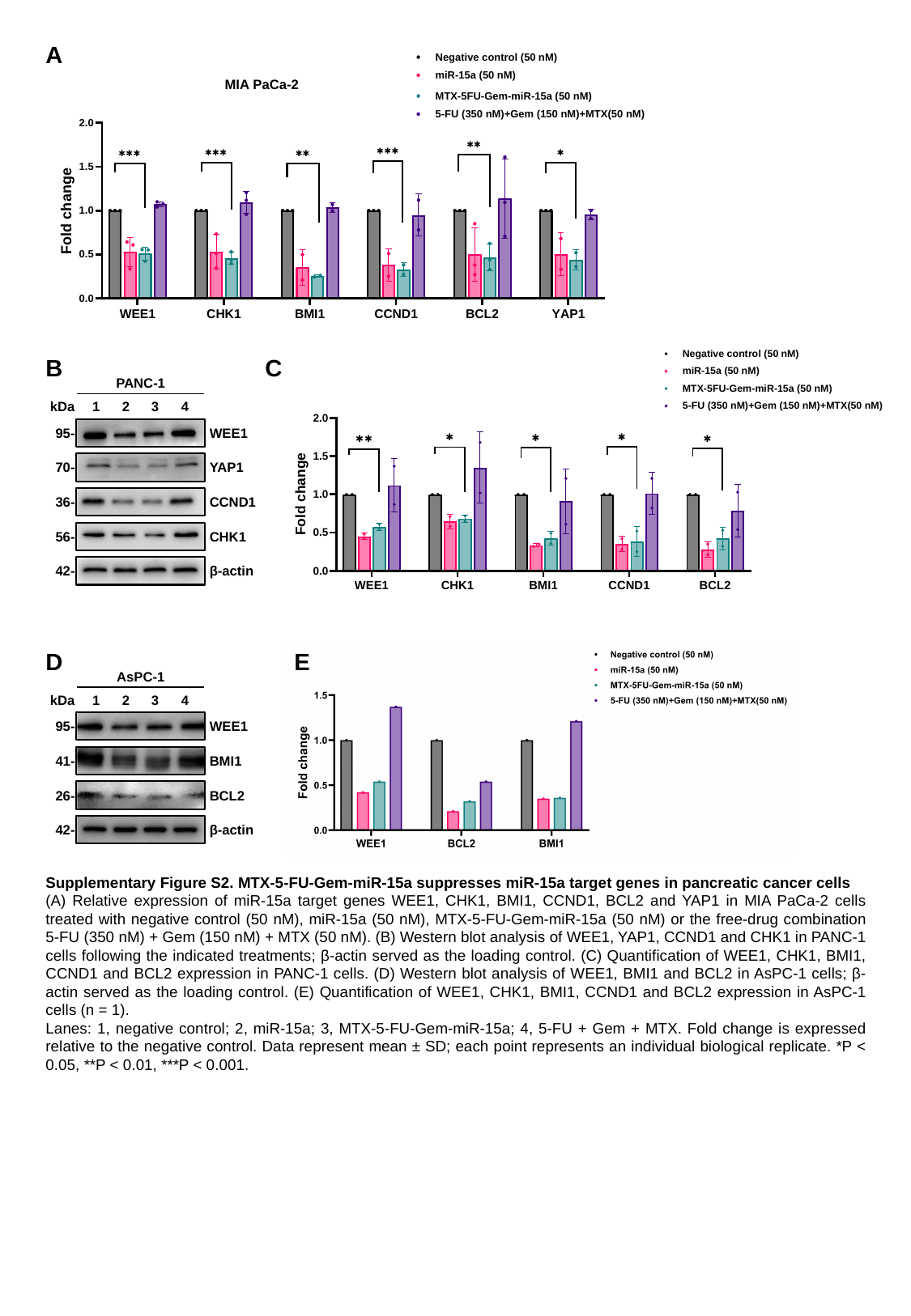

A
MIA PaCa-2
B
C
PANC-1
kDa
1
2
3
4
95-
WEE1
70-
YAP1
36-
CCND1
56-
CHK1
42-
β-actin
D
E
AsPC-1
kDa
1
2
3
4
95-
WEE1
41-
BMI1
26-
BCL2
42-
β-actin
Supplementary Figure S2. MTX-5-FU-Gem-miR-15a suppresses miR-15a target genes in pancreatic cancer cells
(A) Relative expression of miR-15a target genes WEE1, CHK1, BMI1, CCND1, BCL2 and YAP1 in MIA PaCa-2 cells treated with negative control (50 nM), miR-15a (50 nM), MTX-5-FU-Gem-miR-15a (50 nM) or the free-drug combination 5-FU (350 nM) + Gem (150 nM) + MTX (50 nM). (B) Western blot analysis of WEE1, YAP1, CCND1 and CHK1 in PANC-1 cells following the indicated treatments; β-actin served as the loading control. (C) Quantification of WEE1, CHK1, BMI1, CCND1 and BCL2 expression in PANC-1 cells. (D) Western blot analysis of WEE1, BMI1 and BCL2 in AsPC-1 cells; β-actin served as the loading control. (E) Quantification of WEE1, CHK1, BMI1, CCND1 and BCL2 expression in AsPC-1 cells (n = 1).
Lanes: 1, negative control; 2, miR-15a; 3, MTX-5-FU-Gem-miR-15a; 4, 5-FU + Gem + MTX. Fold change is expressed relative to the negative control. Data represent mean ± SD; each point represents an individual biological replicate. *P < 0.05, **P < 0.01, ***P < 0.001.

### Slide 3
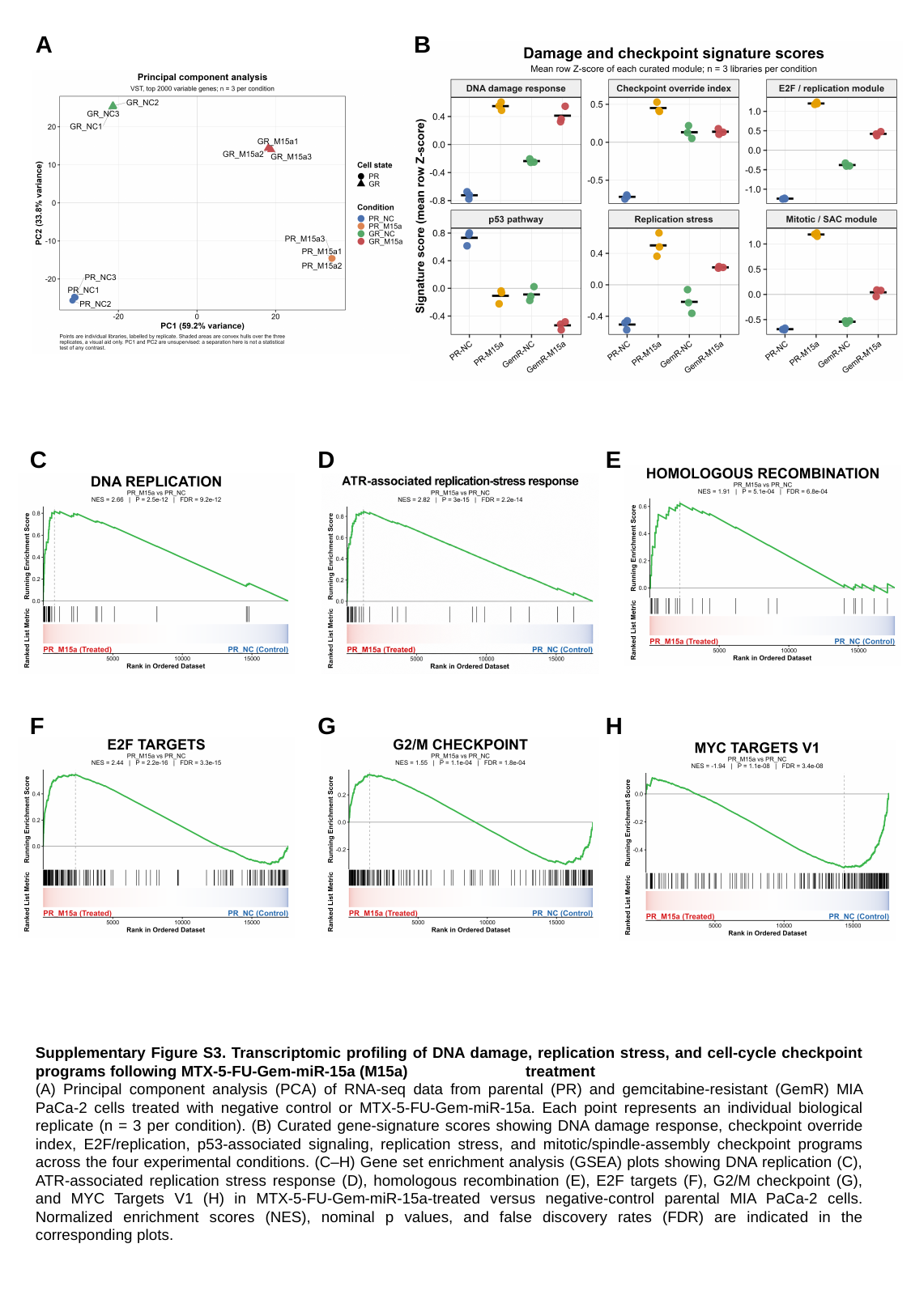

A
B
C
D
E
F
G
H
Supplementary Figure S3. Transcriptomic profiling of DNA damage, replication stress, and cell-cycle checkpoint programs following MTX-5-FU-Gem-miR-15a (M15a)	 treatment
(A) Principal component analysis (PCA) of RNA-seq data from parental (PR) and gemcitabine-resistant (GemR) MIA PaCa-2 cells treated with negative control or MTX-5-FU-Gem-miR-15a. Each point represents an individual biological replicate (n = 3 per condition). (B) Curated gene-signature scores showing DNA damage response, checkpoint override index, E2F/replication, p53-associated signaling, replication stress, and mitotic/spindle-assembly checkpoint programs across the four experimental conditions. (C–H) Gene set enrichment analysis (GSEA) plots showing DNA replication (C), ATR-associated replication stress response (D), homologous recombination (E), E2F targets (F), G2/M checkpoint (G), and MYC Targets V1 (H) in MTX-5-FU-Gem-miR-15a-treated versus negative-control parental MIA PaCa-2 cells. Normalized enrichment scores (NES), nominal p values, and false discovery rates (FDR) are indicated in the corresponding plots.

### Slide 4
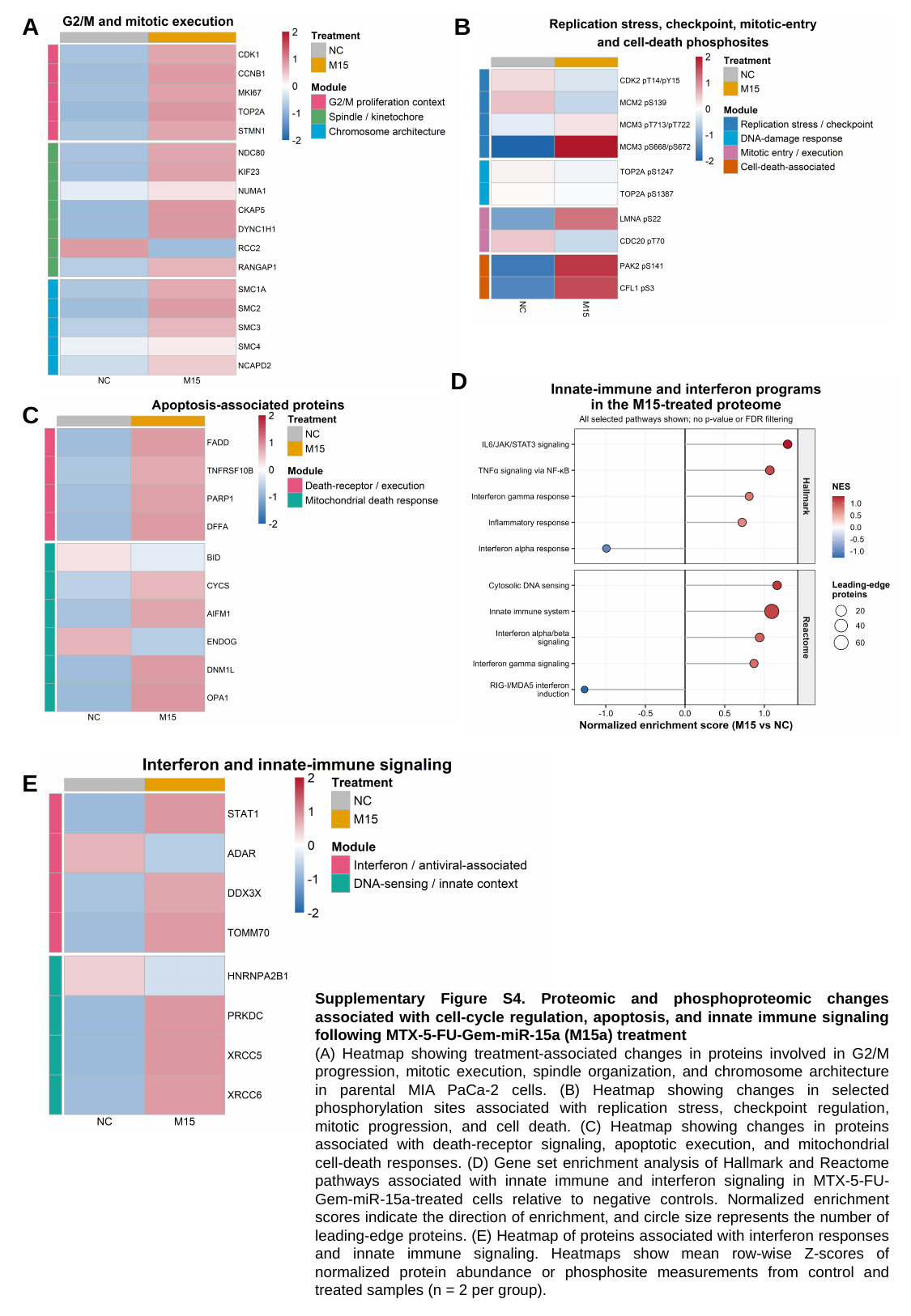

A
B
D
C
E
Supplementary Figure S4. Proteomic and phosphoproteomic changes associated with cell-cycle regulation, apoptosis, and innate immune signaling following MTX-5-FU-Gem-miR-15a (M15a) treatment
(A) Heatmap showing treatment-associated changes in proteins involved in G2/M progression, mitotic execution, spindle organization, and chromosome architecture in parental MIA PaCa-2 cells. (B) Heatmap showing changes in selected phosphorylation sites associated with replication stress, checkpoint regulation, mitotic progression, and cell death. (C) Heatmap showing changes in proteins associated with death-receptor signaling, apoptotic execution, and mitochondrial cell-death responses. (D) Gene set enrichment analysis of Hallmark and Reactome pathways associated with innate immune and interferon signaling in MTX-5-FU-Gem-miR-15a-treated cells relative to negative controls. Normalized enrichment scores indicate the direction of enrichment, and circle size represents the number of leading-edge proteins. (E) Heatmap of proteins associated with interferon responses and innate immune signaling. Heatmaps show mean row-wise Z-scores of normalized protein abundance or phosphosite measurements from control and treated samples (n = 2 per group).

### Slide 5
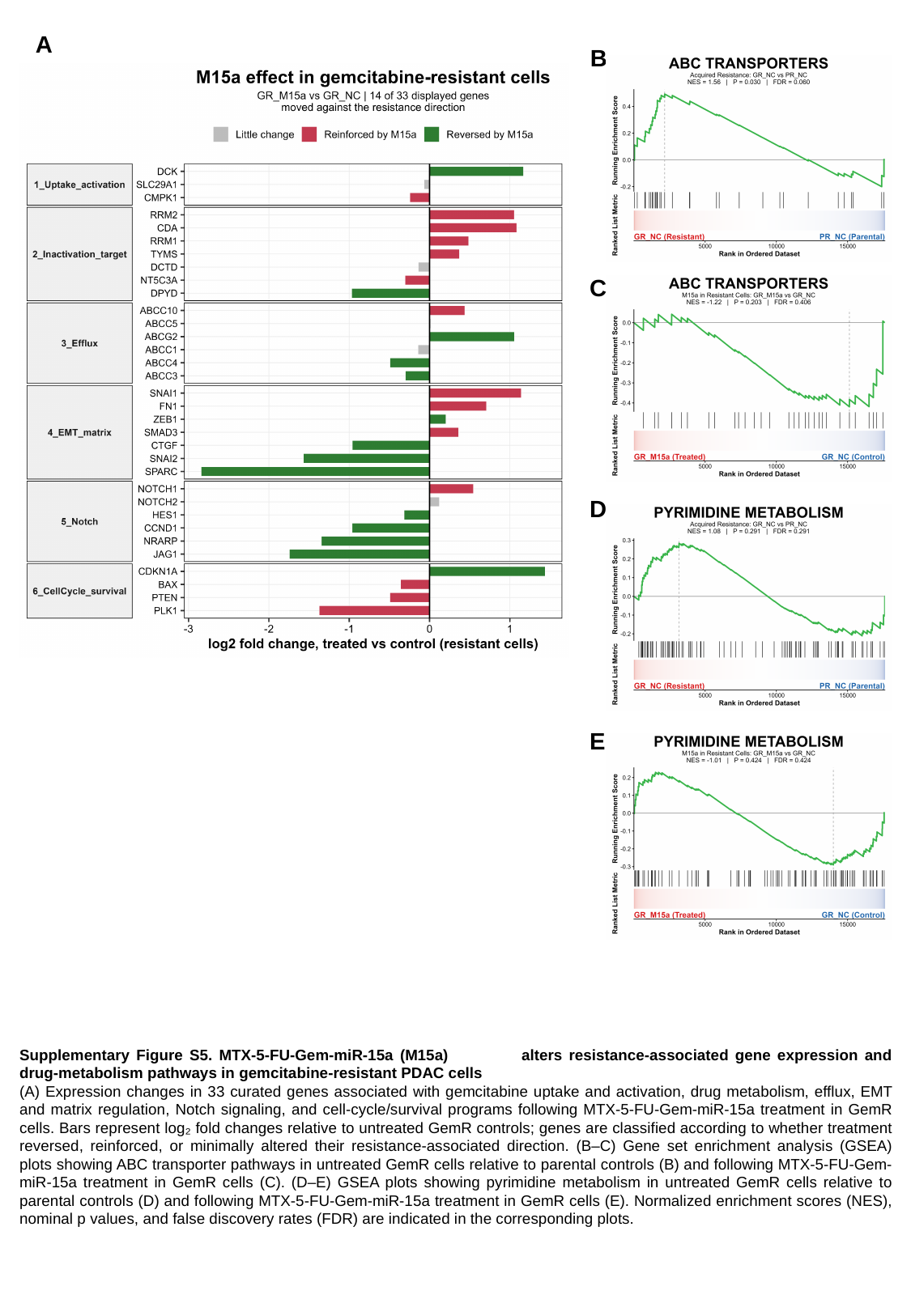

A
B
C
D
E
Supplementary Figure S5. MTX-5-FU-Gem-miR-15a (M15a)	 alters resistance-associated gene expression and drug-metabolism pathways in gemcitabine-resistant PDAC cells
(A) Expression changes in 33 curated genes associated with gemcitabine uptake and activation, drug metabolism, efflux, EMT and matrix regulation, Notch signaling, and cell-cycle/survival programs following MTX-5-FU-Gem-miR-15a treatment in GemR cells. Bars represent log₂ fold changes relative to untreated GemR controls; genes are classified according to whether treatment reversed, reinforced, or minimally altered their resistance-associated direction. (B–C) Gene set enrichment analysis (GSEA) plots showing ABC transporter pathways in untreated GemR cells relative to parental controls (B) and following MTX-5-FU-Gem-miR-15a treatment in GemR cells (C). (D–E) GSEA plots showing pyrimidine metabolism in untreated GemR cells relative to parental controls (D) and following MTX-5-FU-Gem-miR-15a treatment in GemR cells (E). Normalized enrichment scores (NES), nominal p values, and false discovery rates (FDR) are indicated in the corresponding plots.
